# GHB confers neuroprotection by stabilizing the CaMKIIα hub domain

**DOI:** 10.1101/2020.09.28.310474

**Authors:** Ulrike Leurs, Anders B. Klein, Ethan D. McSpadden, Nane Griem-Krey, Sara M. Ø. Solbak, Josh Houlton, Inge S. Villumsen, Stine B. Vogensen, Louise Hamborg, Stine J. Gauger, Line B. Palmelund, Anne Sofie G. Larsen, Mohamed A. Shehata, Christian D. Kelstrup, Jesper V. Olsen, Anders Bach, Robert O. Burnie, D. Steven Kerr, Emma K. Gowing, Selina M. W. Teurlings, Chris C. Chi, Christine L. Gee, Bente Frølund, Birgitte R. Kornum, Geeske M. van Woerden, Rasmus P. Clausen, John Kuriyan, Andrew N. Clarkson, Petrine Wellendorph

**Author notes:** Contributed equally to this manuscript (first author) (Ulrike Leurs, Anders B. Klein, Ethan D. McSpadden). Contributed equally to this manuscript (second author) (Nane Griem-Krey, Sara M. Ø. Solbak).

## Abstract

Ca^2+^/calmodulin-dependent protein kinase II alpha (CaMKIIα) is an abundant neuronal signaling protein involved in synaptic plasticity and memory formation^1,2^. The central hub domain regulates the activity of CaMKIIα by organizing the holoenzyme complex into functional oligomers^3-6^. Recent findings have suggested that the hub is also an allosteric determinant of kinase activity^7^, and is thus an emerging target for therapies to correct CaMKIIα dysregulation^8,9^. However, pharmacological modulation of the hub domain has never been demonstrated. Here we show that stabilization of the CaMKIIα hub domain confers neuroprotection. By combining photoaffinity labeling and chemical proteomics using small molecule analogs of the natural metabolite γ-hydroxybutyrate (GHB)^10^ we reveal that CaMKIIα is the selective target for GHB. We further find that these GHB analogs bind to the hub interior by solving a 2.2 Å crystal structure of CaMKIIα with bound ligand. Using differential scanning fluorimetry, we show that binding of ligands to the hub interior increases the thermal stability of hub oligomers in a concentration-dependent manner. Moreover, we demonstrate the functional significance of this hub stabilization by showing substantial neuroprotective effects in cellular excitotoxicity assays and in a mouse model of cerebral ischemia. Together, our results reveal that CaMKIIα hub stabilization is the mechanism by which GHB provides endogenous neuroprotection and that small-molecule CaMKIIα-selective ligands have therapeutic potential.

## Introduction

The Ca^2+^/calmodulin-dependent protein kinase II alpha (CaMKIIα) is a central mediator of synaptic plasticity governed by its ability to respond to minute fluctuations in Ca^2+^ ^1^. The CaMKIIα holoenzyme is a large protein assembly of 12-14 subunits, each consisting of a kinase domain flexibly linked to the central scaffold hub domain^11^. The hub domain is conserved through evolution^12^ and is endowed with remarkable dynamics^11^. The hub organizes into oligomeric structures^13^, with yet unknown functional importance, however studies clearly highlight the hub as more than an assembling scaffold. The dynamics of the hub domain permits activation-triggered destabilization and release of vertical dimers to enable spreading of activity^5,6^. Furthermore, the hub domain has been reported to interact directly with the kinase domains^4,7,14^ to confer allosteric control of kinase activity^7^. The importance of preserving hub integrity is further evident from a patient hub mutation (p.His477Tyr), which causes defective oligomerization and severe neurodevelopmental defects^15^. Targeting the hub domain thus represents an attractive approach to neuroprotection, especially in cases of CaMKIIα dysregulation, such as excessive activity or hub instability. Thus far, pharmacological modulation of the hub domain has never been demonstrated.

CaMKIIα is activated in a highly cooperative manner, initiated by increases in Ca^2+^, Ca^2+^/CaM binding and autophosphorylation at residue Thr286 in the regulatory segment. In cases of excessive stimuli, such as ischemic brain injury and glutamate-mediated excitotoxicity, a persistent Ca^2+^/CaM-independent autonomous activity is known to persist for hours^9,16,17^ and cause cell death^8^. As a further consequence of Thr286 autophosphorylation, CaMKIIα translocates to the postsynaptic density (PSD) and co-localizes with the NMDA-type glutamate receptor subunit GluN2B^18^. Thus, mechanisms of neuroprotection may either be aimed at inhibition of the kinase or reduction of PSD translocation, both of which have been demonstrated with the cell-permeable CaMKII peptide CN21 (**4**) acting via the kinase domain^19,20^. We here reveal a previously unrecognized endogenous mechanism of compound interaction with a specific binding site in the CaMKIIα hub domain to promote hub stabilization and lasting neuroprotection *in vivo* (Extended Data Fig. 1, schematic summary).

### CaMKIIα is the specific target for GHB in the mammalian brain

γ-Hydroxybutyrate (GHB) is a naturally occurring brain metabolite of γ-aminobutyric acid (GABA) which is reported to be neuroprotective in mammals^21^, however with an elusive mechanism of action. GHB is a weak millimolar agonist at GABA_B_ receptors, mediating sedative and hypothermic effects^22^, but also binds with high affinity to a long-sought elusive binding site, dominant in forebrain regions^23^. This site can be probed selectively with synthetic nanomolar affinity analogs of GHB such as 3-hydroxycyclopent-1-enecarboxylic acid (HOCPCA, **1**)^24^ and 5-hydroxydiclofenac (5-HDC, **2**).

To enable unbiased identification of the long-sought GHB high-affinity binding site, we developed a photolabile diazide-labeled GHB analog, 4-(4-((3-azido-5-(azidomethyl)benzyl)oxy)phenyl)-4-hydroxybutanoate (SBV3, **3**) (*K*_i_ 66 nM), which was employed in an adapted version of photoaffinity labeling, affinity purification and chemical proteomics (Fig. 1, Extended Data Fig. 2A-F)^25^. To this end, **3** was incubated with rat hippocampal membranes in competition with **2**, photoaffinity-labeled, and biotin-ligated. Streptavidin affinity purification followed by LC-MS/MS analysis identified CaMKIIα as the top high-affinity target candidate (R^2^ = 0.81; abundance 7.97×10^9^) (Fig. 1C) and four other less-abundant known CaMKIIα interactors^26^ (Fig. 1D; Extended Data Table 1). Target validation by autoradiography showed complete absence of binding of [^3^H]-**1**^27^ in *Camk2a* -/- mice but not in *Camk2a* +/+, *Camk2b* +/+, and *Camk2b* -/- mice (Fig. 1E). Alpha subtype selectivity was further corroborated by the absence of binding of commercially available radioligands [^3^H]GHB, [^3^H]NCS-382 (Extended Data Fig. 2G-K), and the photoligand **3** to *Camk2a* -/- tissues (Extended Data Fig. 2L-Q). [^3^H]-**1** binding to whole cell homogenate from CaMKIIα-transfected HEK293T cells confirmed saturable binding (Fig. 1F; Extended Data Table 2) and competitive ligand inhibition (Fig. 1G; Extended Data Table 3). No [^3^H]-**1** binding was observed to CaMKIIβ/γ/δ subtypes (Fig. 1H, Extended Data Fig. 3A), thus underscoring the unprecedented CaMKIIα subtype selectivity.

**Fig. 1.**
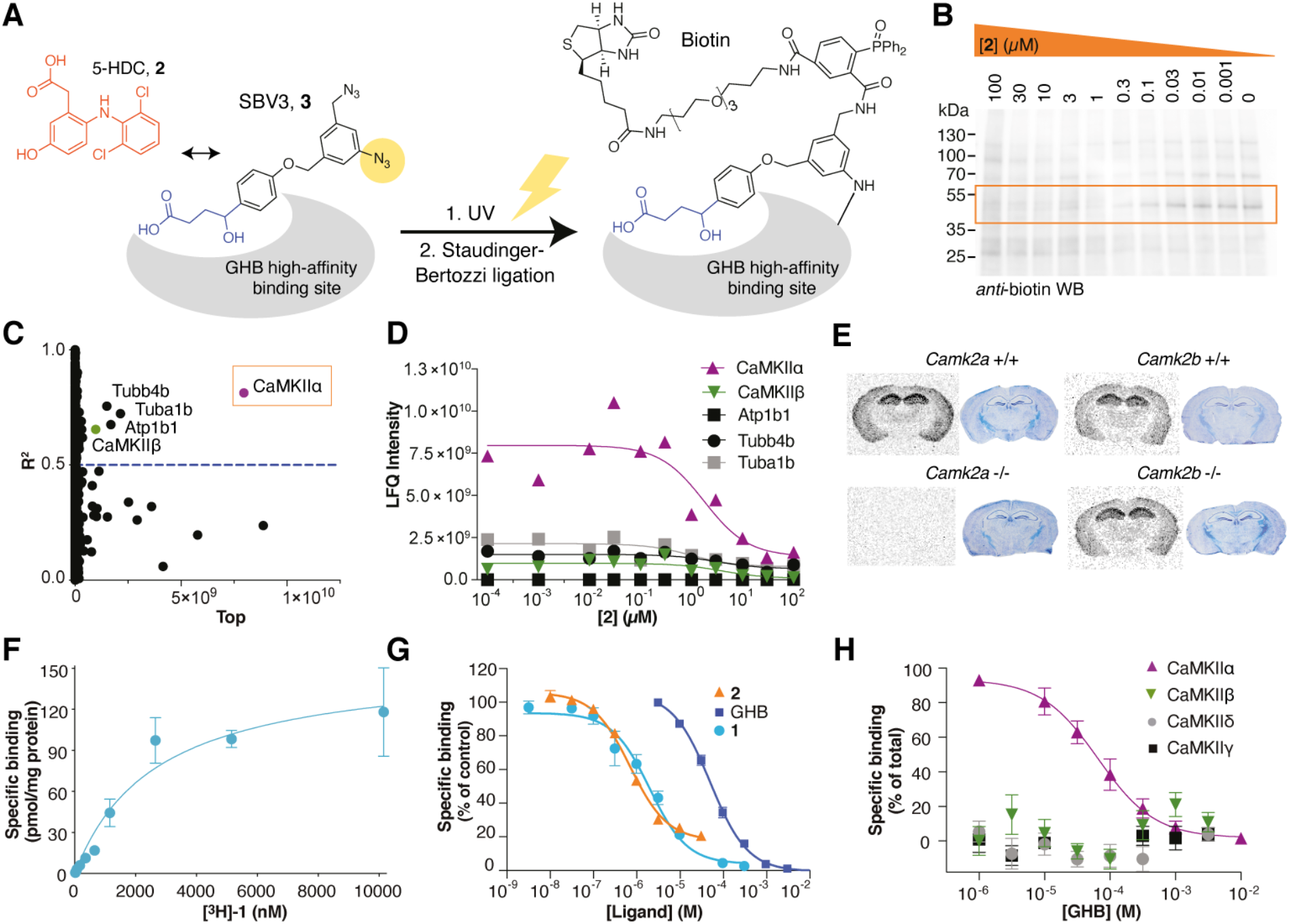
Identification of CaMKIIα as the specific GHB high-affinity target. (A) Approach for target identification using a combination of competitive photoaffinity labeling (PAL) and affinity purification in rat hippocampal homogenate followed by quantitative proteomics. (B) Representative anti-biotin Western blot of hippocampal homogenate after PAL and competition with **2**. (C) Identification of CaMKIIα from LC-MS/MS data as the best hit from non-linear regression analysis for all proteins, and (D) concentration-dependent competition of individual proteins by **2** during PAL. (E) Target validation by [^3^H]-**1** autoradiography using brain slices from *Camk2a and Camk2b* wildtype (+/+) and knockout (-/-) mice (cresyl violet staining for tissue visualization). (F-H) Target validation by [^3^H]-**1** binding to whole cell homogenate from transfected HEK293T cells. (F) [^3^H]-**1** saturation binding to CaMKIIα (*n* = 5), shown is one representative curve; means ± SD). (G) CaMKIIα competition with GHB (*n* = 3), **1** (*n* = 5) and **2** (*n* = 3), pooled data (means ± SEM). (H) Subtype selectivity of [^3^H]-**1** for CaMKIIα cf. CaMKIIβ/γ/δ. Data are pooled (*n* = 3) for each subtype and depicted as specific binding (% of total).

### Structural evidence for a hub domain ligand binding site

Potential ligand binding sites in CaMKIIα include the kinase domain, the regulatory segment and the hub domain (Fig. 2A). From previous crystal structures^3^, we hypothesized that a deep cavity in the hub that contains several positively charged Arg residues is the binding pocket for GHB analogs. Direct binding of GHB analogs to the isolated CaMKIIα hub was demonstrated by surface plasmon resonance (SPR), showing an especially strong binding of **2** (*K*_D_ 0.30 μM) (Fig. 2B, Extended Data Fig. 4A-E). Using a stabilized form of the human hub domain (6x Hub)^12^ we obtained an x-ray crystal structure of a tetradecameric CaMKIIα hub oligomer bound to **2** (2.2 Å resolution) (Fig. 2C-E, Extended Fig. 5A-H). This displayed the direct interactions with Arg residues 433, 453, 469, and His395 in the binding pocket, and also revealed a conformational shift in Trp403 located at the edge of the binding pocket upon binding of **2** (Fig. 2E). These findings were further confirmed using an intrinsic tryptophan fluorescence assay (Fig. 2F, Extended Data Fig. 5J-L), and site-directed mutagenesis of central Arg/His residues into uncharged residues, which consistently disrupted [^3^H]-**1** binding (Fig. 2G, Extended Data Fig. 5J-L). As expected, **1** did not induce a Trp403 flip, likely reflecting its fast kinetics and small size.

**Fig. 2.**
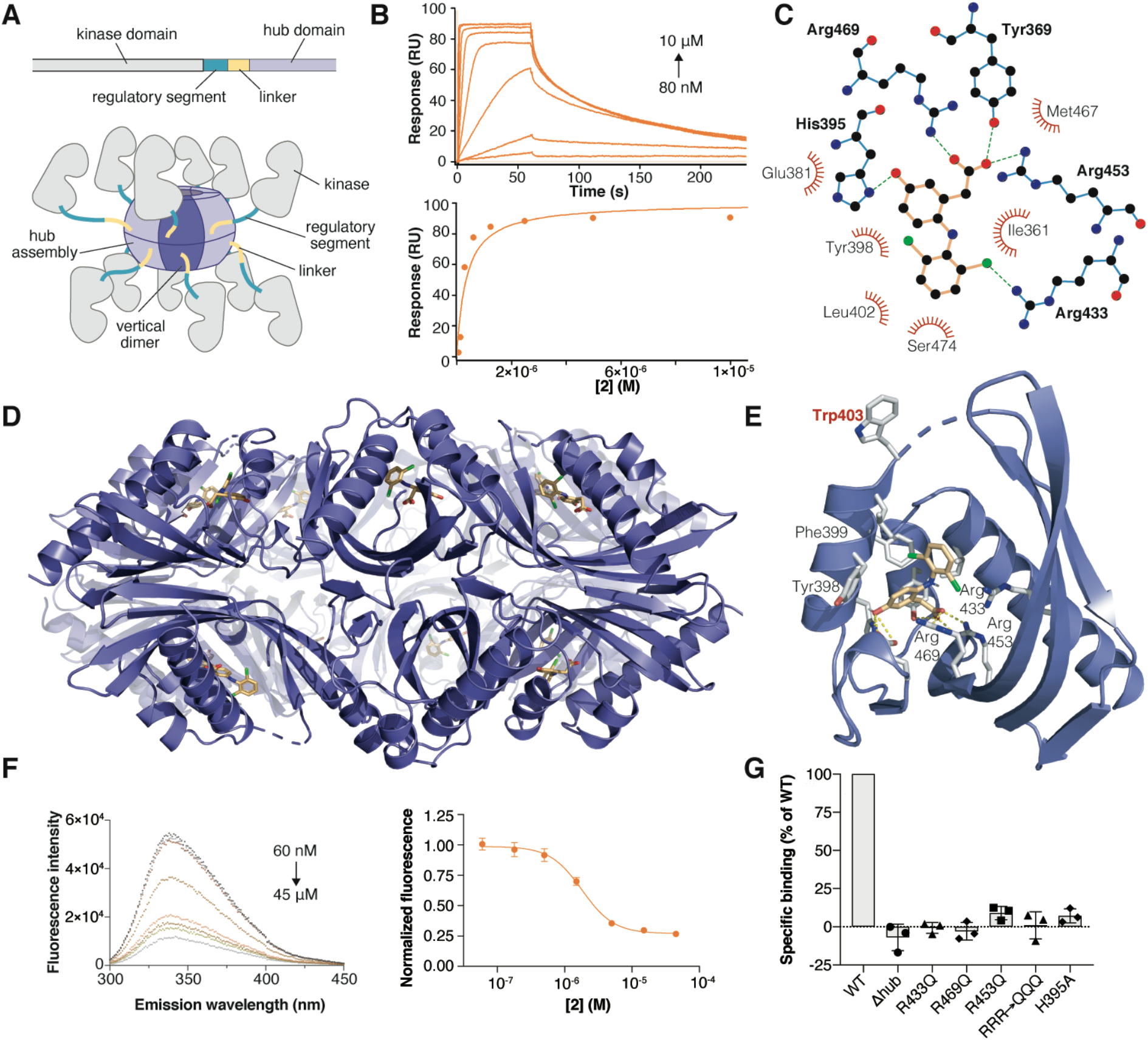
GHB analogs bind the CaMKIIα hub domain. (A) Schematic of a single CaMKIIα subunit comprised of a kinase domain (gray), regulatory segment (green), linker (yellow), and hub domain (lilac). 12-14 hub domains oligomerize into the holoenzyme, shown here in an activated form with kinase domains displaced from the hub. (B) Concentration-dependent binding of **2** to immobilized CaMKIIα 6x Hub measured by SPR (top), and Langmuir binding isotherm (bottom), representative data. (C) Ball and stick model of key binding residues (bold), nearby residues, and hydrogen bonds in green-dashed lines. (D) X-ray crystal structure of **2** bound to the CaMKIIα 6x (14-mer) Hub. (E) Close-up view of a single hub subunit showing the key molecular interactions, displacement (flip) of Trp403 with ligand bound highlighted. (F) Quenching of intrinsic fluorescence caused by Trp403 flip (6x Hub) with increasing concentrations of **2** (*n* = 8), pooled data (means ± SEM). (G) Mutational analysis of key residues in the pocket using [^3^H]-**1** equilibrium binding to CaMKIIα-HEK293T overexpressing cells. Compared to wildtype (WT), binding is completely obliterated in a construct lacking the hub as well as in R433Q, R453Q, R469Q, triple mutant R433/453/468/Q (RRR→QQQ) and H395A mutants (*n* = 3).

### GHB analogs stabilize CaMKIIα hub oligomer formation

Intrigued by GHB analogs binding deep within the hub, we inferred a potential effect of the compounds on hub stability. When testing compounds in a thermal shift assay, pronounced hub-stabilizing effects were observed for GHB, **1** and, most exceedingly, **2**, resulting in respective T_m_ increases of 13, 15 and 29 °C (Fig. 3A, Extended Data Fig. 4F-H) that correlate with the relative affinity rank order of the compounds at pH 6 (Extended Data Table 3). Such a stabilization of the hub oligomer seems an attractive mode of action to counter-balance excessive CaMKIIα activity, such as in cases of excitotoxicity or ischemia. In line with the hub site of action of **1**, it did not affect Thr286 autophosphorylation in cultured cortical neurons (Fig. 3B; Extended Data Fig. 6A-H), and did not affect classical LTP induction in hippocampal slices (Extended Data Fig. 7A-D). Neither did GHB, **1** or **2** affect substrate phosphorylation of syntide-II (Extended Data Fig. 7E).

**Fig. 3.**
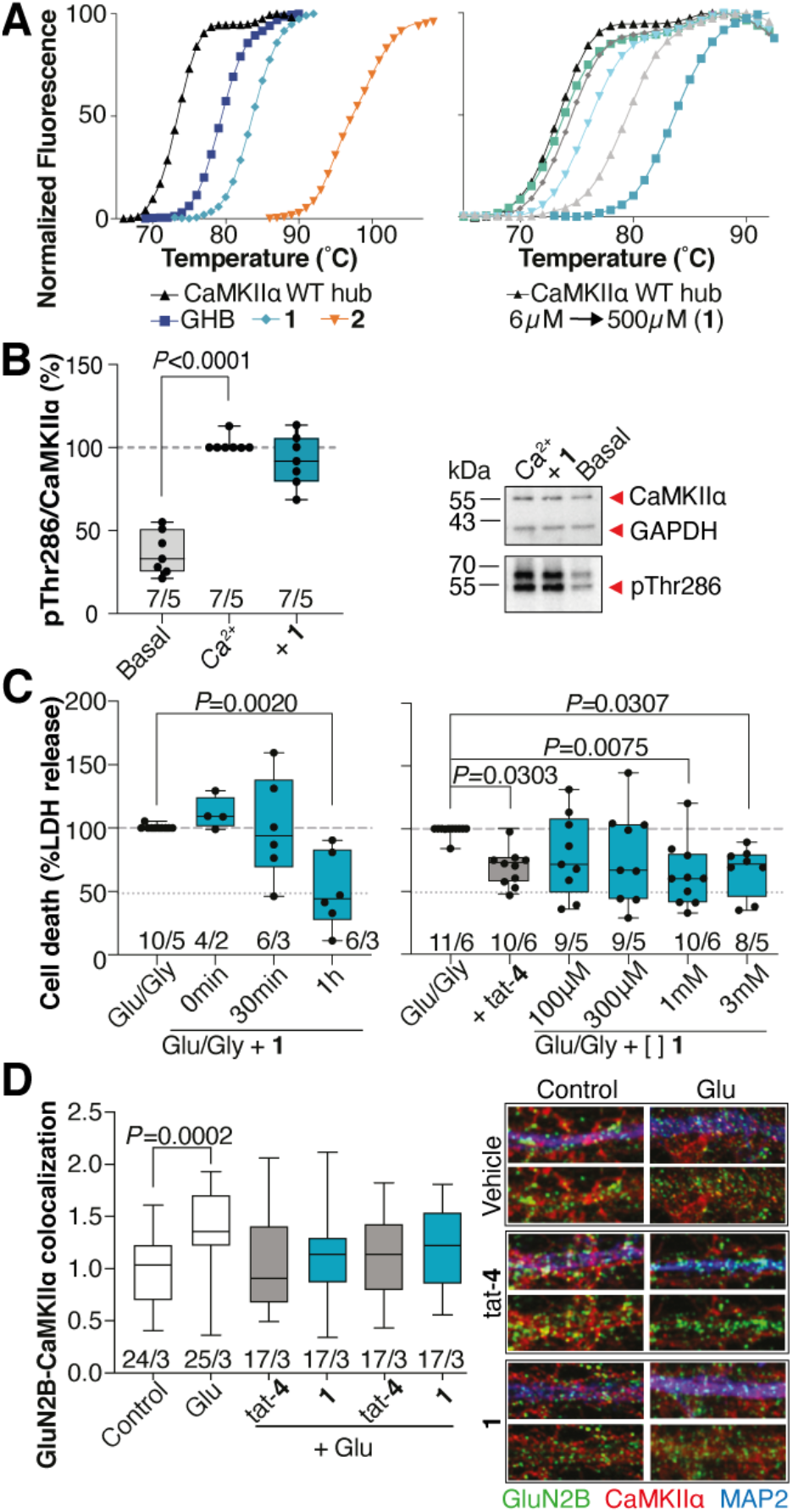
Functional effects of GHB analogs. (A) Right-shifted thermal shift assay melting curves of CaMKIIα WT hub upon binding of GHB, **1** and **2** (*left*) and **1** concentration-dependence (right), representative data. (B) No effect of **1** on Ca^2+^-stimulated Thr286 phosphorylation. Shown is quantification of mean band intensities of Ca^2+^-stimulated pThr286 levels (*left*) normalized to total CaMKIIα expression of cultured cortical neurons (DIV 18-20) incubated with 50-100 μM Ca^2+^ alone or together with 3 mM of **1** for 1 h and representative Western blots (*right*). GAPDH was used as loading control (C) Time-(*left*) and concentration-dependent effects of **1** (*right*) on cell survival at 24 h in cultured cortical neurons (DIV 16-18) stimulated with 100-200/20 μM Glu/Gly for 1 h (tat-**4** as control). Cell death was normalized to maximum cell death as measured by LDH release. (One-way ANOVA, post-hoc Dunnett’s test). (D) Quantification of GluN2B-CaMKIIα co-localization in hippocampal neurons (DIV 14-19) exposed to Glu 400 μM for 2 min and immediately fixed (*left*) and representative immunostained images (*righ*t). *For B-D:* Number in bar diagrams indicates number of experiments/individual cultures. Box plots (boxes, 25–75%; whiskers, minimum and maximum; lines, median). (One-way ANOVA, post-hoc Dunnett’s test).

### CaMKIIα hub ligands are cellular protectants

To study the mechanistic effects of CaMKIIα oligomeric stabilization we selected the highly CaMKIIα hub-selective ligand **1** for *in vitro* studies in neuronal cultures. Where GHB is known to cause hypothermia and sedation via metabotropic GABA_B_ receptors^22^, **1** has no affinity for GABA_B_ receptors^24^, nor does it induce hypothermia which could by itself lead to neuroprotection (Extended Data Fig. 7F). In cultured neurons, we initially examined effects of **1** by exposure to glutamate excitotoxicity and cell death measurements in a lactate dehydrogenase (LDH) cell viability assay (Extended Data Fig. 6I)^28^. When applying **1** (1 mM) at different times post-noxius stimulus, a significant concentration-dependent protective effect was observed at 1 hour (Fig. 3C). In keeping with a CaMKIIα-dependent neuroprotective action, we observed an inhibitory effect of **1** (similar to tat-**4**) on GluN2B-CaMKIIα co-localization in glutamate-stimulated hippocampal dendritic spines (Fig. 3D; Extended Data Fig. 8)^29^.

### CaMKIIα hub ligands afford lasting neuroprotection *in vivo*

We next investigated treatment effects of **1** in a mouse model of photothrombotic focal ischemic stroke^30^. Similar to other models of stroke, we confirmed a CaMKIIα-relevant pathogenesis as evidenced by an elevation in pThr286 levels 3-12 hours post-injury (Extended Data Fig. 9A-C). Both GHB and **1** significantly reduced the infarct volume when given 30 min post-stroke (Extended Data Fig. 9D). Furthermore, when **1** (175 mg/kg) was administered at 3, 6 and 12 hours post-photothrombotic stroke (Fig. 4A), it resulted in a significant decrease in infarct volume by ~40-50% measured 7 days post-stroke (Fig. 4B). This was accompanied by improvements in motor coordination measured in paw placement on the grid-walking task (Fig. 4C), and in forelimb asymmetry in the cylinder test (Extended Data Fig. 9E-F). These effects were equally attained with a lower dose of compound (90 mg/kg) (Extended Data Fig. 9G-H). Of clinical relevance, **1** (175 mg/kg) was also found to reduce infarct sizes in aged female mice (Fig. 4D). Finally, we investigated the effect of **1** on the functional consequences of a motor cortex stroke in relation to interhemispheric transfer and integration of sensorimotor and cognitive information via axons through the corpus callosum 14 days post-stroke^31^. Recording of compound action potentials (CAPs) (Fig 4E-G) showed an overall significant increase in CAPs with treatment (~50-60% of control) (Fig. 4G). The impairments in axon function were further supported by a compound-mediated reversal of the stroke-induced decrease in transport of the neuroanatomical tracer biotinylated dextran amine (BDA) (Extended Data Fig. 9I-J).

**Fig. 4.**
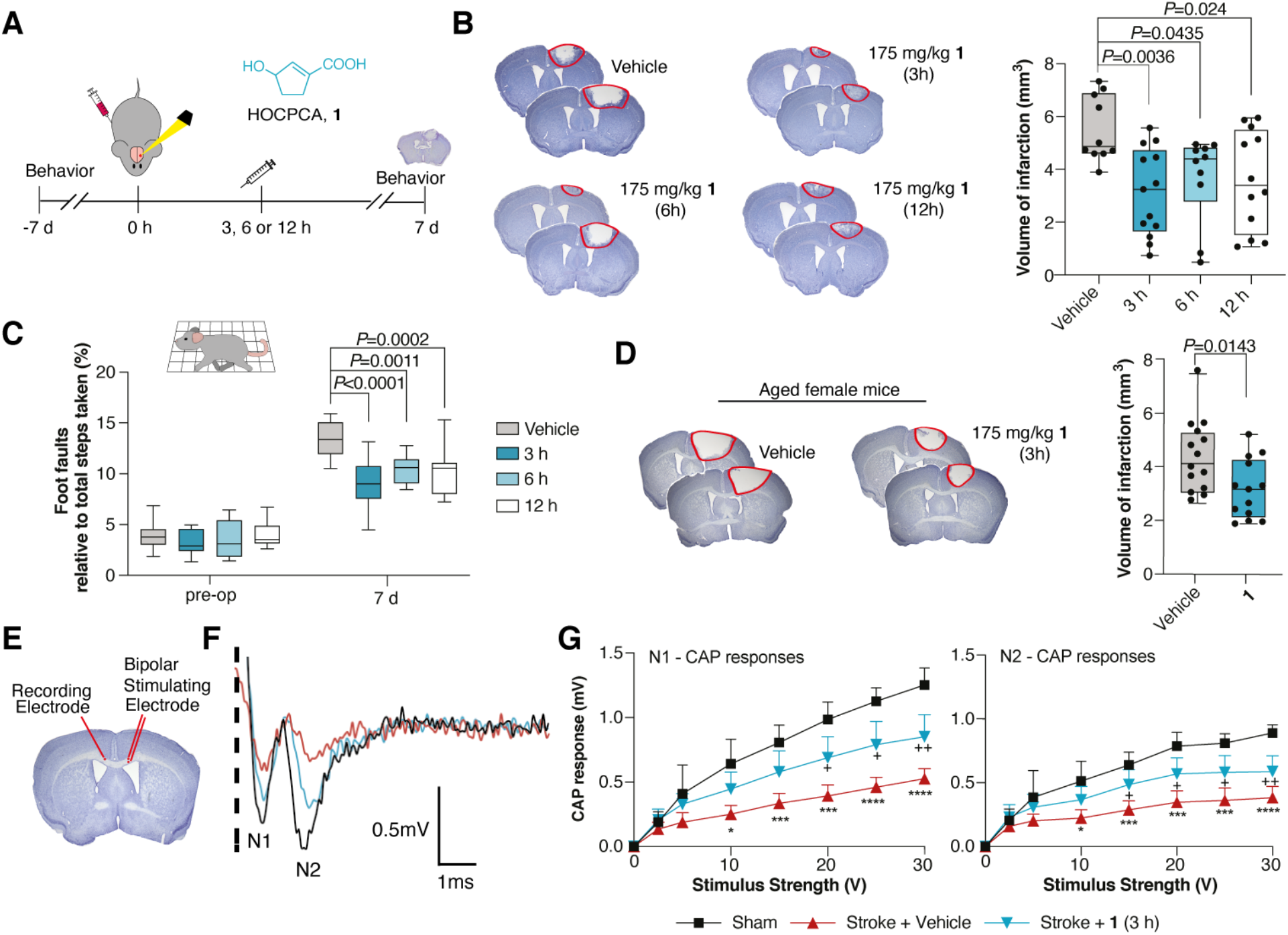
Neuroprotective effects of compound 1 *in vivo*. (A) Time line for testing of the GHB analog **1**. (B) Effect on infarct size after treatment with a single dose of compound **1** at 3, 6 or 12 h post-photothrombosis (One-way ANOVA, post-hoc Dunnett’s test), representative cresyl violet staining with doses and n’s indicated (*left*) and quantification of stroke volumes (*right*). (C) Functional recovery assessed in grid-walking task (Pre-op, pre-operation; Two-way ANOVA (time, treatment), post-hoc Dunnett’s test). (D) Treatment of aged female mice (20-24 months) with **1** (Two-tailed Student’s *t*-test), representation as for (B). (*E-G*) Axonal function assessed by electrophysiological recording of compound action potentials (CAPs) 14 days post-stroke. Mice were treated with a single dose of 175 mg/kg of **1** at 3 h post-stroke (blue) cf. vehicle (red) and sham (black). (F) Representative recording of CAPs showing negative peak for myelinated (N1) and unmyelinated axons (N2). (G) Amplitudes of CAP peaks for N1 (*right*) and N2 (*left*). (Means ± SD, Two-way ANOVA (stimulus strength, treatment), post-hoc Tukey’s test, +*P*<0.05, ++*P*<0.01, sham compared to stroke + **1**, \**P*<0.05, \*\*\**P*<0.001, \*\*\*\**P*<0.0001 sham compared to stroke + vehicle.) Box plots (boxes, 25–75%; whiskers, minimum and maximum; lines, median).

## Discussion

The role of the CaMKIIα hub as a determinant for holoenzyme assembly and structural integrity is well-established^3,6,7^, yet its functional importance has been less appreciated. This study identifies a novel binding site in the CaMKIIα hub domain that, incidentally, corresponds to the enigmatic high-affinity site for GHB in the mammalian brain. Its unequivocal identification now offers new mechanistic understanding of GHB physiological relevance and efficacy in neuroprotection^21,32^. Though GHB itself also affects GABA_B_ receptors^22^, hub stabilization may well explain its already applied pharmacological efficacy in conditions like alcoholism and narcolepsy^33^. Compounds with a more specific mode of action (like **1**) are therefore advantageous in several states of CaMKII dysregulation.

The CaMKIIα hub-structure presented here is the first with a bound ligand. The central Arg residues 453 and 469 in the pocket make important interactions with the carboxylic acid moiety, as previously suggested^3,12^. The fact that the co-crystal with **2** is a tetradecamer further infers that this oligomeric state is stabilized by GHB and analogs. The Trp403 displacement upon binding shows a new aspect of hub molecular dynamics. As this residue is conserved only to CaMKIIα, it may be an important determinant of the subunit selectivity filter for this type of compounds. Interestingly, Trp403 was recently reported as a part of a loop mediating kinase docking in CaMKIIα lacking a linker region^7^, suggesting that a conformational change in this part of the hub may also influence allosteric control of holoenzyme activity.

Our structural and molecular data suggest a prototypical action of small-molecule hub ligands, such as GHB analogs, in providing pharmacological compensation of CaMKIIα overactivity by stabilization of oligomer formation. In relation to cerebral ischemia, this mechanism appears to be of particular relevance several hours post-insult, where CaMKIIα is known to have excessive autonomous activity^9,17,20^ and translocate to the PSD^18^. Given the limited treatment options and general poor prognosis for functional recovery after cerebral ischemia^34^, targeting the lasting activation of CaMKIIα offers an extended time window for therapeutic rescue of neuronal tissue at risk. Overall, our elucidations present a new principle for regulating CaMKIIα function by inhibition of hub dynamics.

## Supporting information

Mass spec data

## Acknowledgements

We thank members of all labs for their support. We appreciate insightful comments from Ype Elgersma, Howard Schulman, and expert advice from Bernhard Küster and Mike Gibson. A special thanks to Kresten Lindorff-Larsen and Nils Ole Dalby for invaluable input. Mehrnoush Aghadavoud Jolfei is kindly acknowledged for help with the LTP experiments, Ales Marek for providing [^3^H]-**1** radioligand, and Anders Skov Kristensen for providing access to Safire^2^ plate reader. For x-ray crystallography data we thank James Holton and George Meigs at the Advanced Light Source Beamline 8.3.1 for assistance with data collection. Beamline 8.3.1 at the Advanced Light Source is operated by the University of California Office of the President, Multicampus Research Programs and Initiatives grant MR-15-328599, the National Institutes of Health (R01 GM124149 and P30 GM124169), Plexxikon Inc. and the Integrated Diffraction Analysis Technologies program of the US Department of Energy Office of Biological and Environmental Research. The Advanced Light Source (Berkeley, CA) is a national user facility operated by Lawrence Berkeley National Laboratory on behalf of the US Department of Energy under contract number DE-AC02-05CH11231, Office of Basic Energy Sciences.

## Author contributions

U.L., R.P.C. and P.W. conceptualized the original idea, E.D.M. and J.K. proposed the hub binding site. A.B.K., S.M.Ø.S. B.R.K., G.M.v.W., A.N.C. and P.W. further developed the concept. U.L., A.B.K., E.D.M., N.G.-K., J.H., I.S.V., S.B.V., S.M.Ø.S., L.H., S.J.G., L.B.P., A.S.G.L., C.D.K., C.C.C., C.L.G., R.O.B., S.K., E.K.G., S.M.W.T., C.G., M.A.S., G.M.v.W., A.N.C. and P.W. performed the experiments and analyzed the data. B.F. and R.P.C. developed and provided essential compounds. C.D.K. and J.V.O. provided mass spectrometry resources. U.L., A.B.K., N.G.-K., A.N.C., and P.W. wrote the manuscript with input from all authors. N.G.-K. and P.W. performed data visualization. P.W. supervised and administered the project overall.

## Methods section

### Materials

#### Compounds and radioligands

GHB (γ-hydroxybutyrate) sodium salt, 5-HDC (5-hydroxydiclofenac, **2**) and KN-93 (**6**) were purchased from Sigma-Aldrich (St. Louis, MO, USA). HOCPCA (3-hydroxycyclopent-1-enecarboxylic acid, **1**) sodium salt was synthesized as described previously^35^ and is available in small amounts by request. The engineered photoligand, SBV-3 (**3**), was synthesized as described. Purity of ligands were at least 95%. [^3^H]NCS-382 ([^3^H]-**5**, spec. activity 20 Ci/mmol, #ART-1114) and [^3^H]GHB (20 Ci/mmol, #ART-0365) were purchased from Biotrend (Köln, Germany). [^3^H]HOCPCA ([^3^H]-**1**, 28.6 Ci/mmol) was prepared as previously described ^35^. Peptides used for control experiments (CaM binding element peptide (CBEP, **7**), LKKFNARRKLKGAILTTMLA; syntide-II, PLARTLSVAGLPGKK; CN21, KRPPKLGQIGRSKRVVIEDDR (**4**); TatCN21, GRKKRRQRRRKRPPKLGQIGRSKRVVIEDDR (tat-**4**)) were obtained from Genscript Biotech (Leiden, Netherlands). Peptides were acetylated at the C-terminus, amidated at the N-terminus, purified by HPLC, and subjected to standard trifluoroacetic acid (TFA) removal (as TFA was found to interfere with ligand binding). Peptide **7** corresponds to amino acids 290-309 of the autoinhibitory segment of human CaMKIIα^36^. Standard buffers were made using high-grade reagents and solvents purchased from Sigma-Aldrich unless otherwise noted.

#### Antibodies and Western blotting materials

The following primary antibodies were used: streptavidin-HRP (#434323, RRID:AB_2619743; Invitrogen; validated in this study Extended Data Fig. 2C), CaMKIIα (#NB100-1983, RRID:AB_10001339; mouse monoclonal IgG, clone 6g9, Novus Biologicals; validated in this study Extended Data Fig. 2N), CaMKIIβ (#12716, RRID:AB_2713889; mouse monoclonal IgG2b, clone CB-beta-1, Invitrogen; validated in this study, Extended Data Fig. 2P), phospho-CaMKII (Thr286) (# AP12716b, RRID:AB_10820669; rabbit monoclonal IgG, clone D21E4, Cell Signaling Technology; validated in this study, Extended Data Fig. 6F), GluN2B (#21920-1-AP, RRID:AB_11232223; rabbit polyclonal IgG, Proteintech; validated by the company (https://www.ptglab.com/products/GRIN2B-Antibody-21920-1-AP.htm#validation), MAP2 (#188 004, RRID:AB_2138181; guinea pig polyclonal antiserum, Synaptic Systems; validated by the company (https://www.sysy.com/products/map2/facts-188004.php)), GAPDH (#NB300-221, RRID:AB_10077627; mouse monoclonal IgG, clone 1D4, Novus Biologicals), Myc tag (#MA1-21316, RRID:AB_558473; mouse monoclonal IgG1, Invitrogen), Na^+^/K^+^-ATPase (#ab76020, RRID:AB_1310695; rabbit monoclonal, clone EP1845Y, Abcam). Secondary antibodies used were HRP-rabbit anti-mouse (#P0161, RRID:AB_2687969; polyclonal Ig fraction, Agilent), HRP-goat anti-rabbit (#P0448, RRID:AB_2617138; polyclonal, Agilent), Goat anti-mouse Alexa Fluor Plus 647 (#A32728, RRID:AB_2633277; polyclonal IgG, Invitrogen), polyclonal IgG, Jackson ImmunoResearch). Final antibody dilutions used, re-using of antibody solutions and their storage are stated in the relevant subsections.

For Western blotting, a ‘1% protease/phosphatase inhibitor cocktail’ was customarily used, consisting of 1% cOmplete™ protease inhibitor cocktail (Roche Diagnostics), 1% phosphatase inhibitor cocktail 2 (#P5726, Sigma-Aldrich) and 1% phosphatase inhibitor cocktail 3 (#P0044, Sigma-Aldrich). For protein determination, either the Bradford assay (#5000006, Bio-Rad Protein Assay Dye Reagent Concentrate; Bio-Rad Laboratories, Copenhagen, Denmark) or the Pierce™ BCA Protein Assay Kit (#23227, Thermo Fisher Scientific, West Palm Beach, FL, USA) were used according to the manufacturer’s instructions, as specified.

#### Plasmids and mutants

Plasmids were all rat: pCMV6-CaMKIIα-Myc-DDK, (#RR201121), pCMV6-CaMKIIγ-Myc-DDK (#RR207416), pCMV6-CaMKIIδ-Myc-DDK (#RR209882), all from Addgene, pCAGG-CaMKIIβ-pPGK-tdTOMATO ^37^. Mutations R433Q, R453Q, R469Q, H395A and the triple mutant R433/453/469Q in CaMKIIα in pCMV6-Myc-DDK were generated and sequence-verified by GenScript. Heterologous expression of CaMKIIα hub domains (wildtype; WT and 6x mutated hub^38^) in *E. coli* was done using a pSKB2 plasmid with kanamycin resistance.

#### Cell lines and primary cultures

The human embryonic kidney (HEK) 293T cell line was purchased from ATCC (293T-ATCC; #CRL-3216; authenticated to be mycoplasma-free). Primary cortical neurons were prepared from E16-E18 embryos and hippocampal neurons from E16.5 embryos from time-mated female C57BL6/*J*Rj female mice (Janvier Laboratories, Le Genest-Saint-Isle, France) or from genetically modified *Camk2a* -/- or +/+ mice as described).

#### Animals and mouse breeding (general)

Specific details pertaining to age, weight and sex of animals used are given in the relevant sections. All breeding protocols and animal experiments were approved by the respective University animal ethics committees following national and international guidelines.

Rats for *in vitro* binding studies were Sprague Dawley obtained from commercial breeders (Janvier) (RRID: RGD_7246927). Mice for LTP recordings were C57BL/6JOlaHsd mice from inhouse breeding. Mice for *in vitro* studies (homogenate binding and autoradiography) were *Camk2a* (*Camk2a*^tm3Sva^, MGI:2389262) or *Camk2b* knockouts (-/-) or corresponding litter mates (+/+) backcrossed in the C57BL/6*J* background. Generation of *Camk2b* knockout mouse lines has been described elsewhere^39^. *Camk2* lines were bred as heterozygotes to generate +/+ and -/- litter mates, except for neuronal cultures where homozygous breeding, between either +/+ or -/- mice was also used. Genotyping was performed by a technician blinded to further experiments. Mice for in vivo experiments are further specified in the relevant sections.

### Chemistry

#### Synthesis of SBV3 precursor

*5-(4-((3-Azido-5-(azidomethyl)benzyl)oxy)phenyl)dihydrofuran-2(3H)-one* SBV3-43 was synthesized from precursors 3-azido-5-(azidomethyl)benzyl methanesulfonate and 5-(4-hydroxyphenyl)dihydrofuran-2(3*H*)-one ^40^. 3-azido-5-(azidomethyl)benzylmethanesulfonate (681 mg, 2.41 mmol) and 5-(4-hydroxyphenyl)dihydrofuran-2(3*H*)-one (430 mg, 2.41 mmol) were dissolved in DMF (20 mL) before K_2_CO_3_ (500 mg, 3.62 mmol) was added and the resulting solution was stirred at 70 °C overnight. The thick, dark solution was poured into HCl (aq.) (0.1 M, 75 mL) and the aqueous phase was extracted with EtOAc (3 x 75 mL), washed with water (2 x 50 mL) and brine (20 mL), dried (MgSO_4_) and the solvent was removed *in vacuo*. Column chromatography (2:1 Heptane/EtOAc) yielded SBV3 precursor as a yellow oil (502 mg, 1.38 mmol, 57%). ^1^H NMR (CDCl_3_)): δ 2.14–2.22 (m, H), 2.56–2.68 (m, 3H), 4.35 (s,2H), 5.04 (s, 2H), 5.43–5.46 (m, 1H), 6.91–6.94 (m,1H), 6.97 (d, J = 8.8, 2H), 7.06–7.08 (m, 1H), 7.14–7.16 (m, 1H), 7.27 (d, J = 8.4 Hz, 2H). ^13^C NMR (CDCl_3_)): δ 29.27, 30.83, 54.11, 69.15, 81.39, 115.06, 117.54, 118.06, 123.22, 127.19, 131.88, 137.96, 139.62, 158.56, 177.20. LC-MS: MH^+^: 365.1.

#### Synthesis of SBV3 (3)

*Lithium 4-(4-((3-azido-5-(azidomethyl)benzyl)oxy)phenyl)-4-hydroxybutanoate* 5-(4-((3-Azido-5-(azidomethyl)benzyl)oxy)phenyl)dihydrofuran-2(3*H*)-one (50 mg, 0.137 mmol) was dissolved in THF (100 μL) and LiOH (aq) (206 μL, 2N, 0.41 mmol) was added. The orange solution was stirred for 2 h at room temperature. Water was added (10 mL) and the aqueous phase washed with diethyl ether (2 x 3 mL). The aqueous phase was evaporated to give the product as a yellow solid (75 mg, 100%). ^1^H NMR (D_2_O, dioxane): δ 1.81–1.96 (m, 2H), 2.00–2.11 (m, 2H), 4.22–4.25 (m, 2H), 4.49–4.52 (m, 1H), 4.90–4.95 (m, 2H), 6.85–6.92 (m, 3H), 6.96–6.99 (m, 1H), 7.06–7.10 (m, 1H). ^13^C NMR (D_2_O, dioxane): δ 34.12, 34.52, 53.66, 69.34, 73.28, 115.16, 117.79, 118.32, 123.96, 127.68, 136.84, 138.08, 139.43, 140.82, 157.22, 167.36, 171.15, 182.80. LC-MS (M-N_2_-OH)H^+^ : 337.20.

### Ligand characterization using rat brain homogenate

#### Homogenate preparation

Rat cerebral cortical P2 synaptosomally-enriched membranes (referred to as ‘homogenate’) were prepared from healthy adult male (250**–**300 g) Sprague Dawley rats, as described earlier^41^. Typically, 20-23 rats were decapitated and tissue pooled for homogenate preparations. Care was taken to collect brains right after decapitation. The final pellet (5 x volume/weight) was washed 3-4 times in binding buffer (50 mM KH_2_PO_4_ buffer, pH 6.0) by a centrifugation-resuspension procedure using a SS-34 fixed angle rotor in a high-speed Sorvall centrifuge (48,000 x *g*) and aliquoted and stored at -20 °C. On the day of the assay, homogenates were quickly thawed by shaking in binding buffer, pelleted and resuspended in binding buffer to approx. 5 mg/mL. Protein concentrations were determined using the Bradford method. Homogenates were kept on ice until use.

#### Equilibrium binding assays (Extended Data Fig. 2A-B)

Inhibitory affinities (*K*_I_ values) of **2** and **3** were determined using the well-established rat cortical homogenate competition assays employing either [^3^H]-**5** (16 nM) or [^3^H]-**1** (5 nM) exactly as previously described^35,41^. Due to a limited amount of compound, **3** was only determined in the [^3^H]-**5** assay. In brief, homogenate amounting to 25–40 μg total protein per 96-well was mixed with radioligand and test compound in a total volume of 200 μL. Nonspecific binding (NSB) was determined with 1 mM GHB. After 1 h incubations at 0-4 °C, reactions were terminated by rapid filtration through GF/C unifilters (PerkinElmer, Waltham, MA, USA) using a cell harvester (Packard), washed quickly three times with ice-cold binding buffer, dried, and added scintillation liquid MicroScint-0 (PerkinElmer) before counting plates for radioactivity (3 min per well) in a TopCount NXT Microplate Scintillation counter (PerkinElmer). To permit *K*_I_ calculations, *K*_D_ values as well as actual radioligand concentrations from each experiment were used. Whereas the *K*_D_ value for [^3^H]-**5** (430 nM) has been reported previously ^41^, the *K*_D_ value for [^3^H]-**1** (259 nM) was determined by a saturation experiment using radioligand concentrations ranging from 10-15,000 nM and isotope dilutions at concentrations above 500 nM, 5 mM GHB for NSB and otherwise using the same protocol as for competition binding.

#### Data analysis

All homogenate binding assays were performed in technical triplicates using pooled membranes. Data are visualized as pooled data of at least three independent experiments (means ± SEM). Competitive inhibition curves were analyzed by the one-site model by nonlinear regression using:

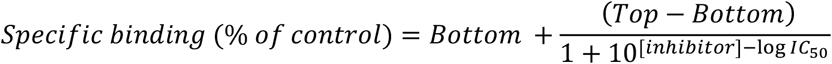

where bottom and top represent the upper and lower plateaus, respectively, of the sigmoidal binding curve and where specific binding was corrected for NSB and normalized to the total signal. IC_50_ values were converted to *K*_I_ values using the Cheng-Prusoff equation:

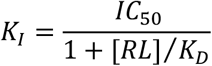

where [RL] is the added concentration of radioligand and *K*_D_ is the dissociation constant determined from saturation experiments.

Saturation data were fitted to a one-site model by non-linear regression to determine *K*_D_ and B_max_:

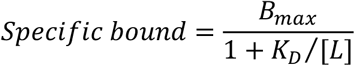

Specific bound was corrected for NSB determined from each radioligand concentration. For concentrations based on isotope dilution, NSB was extrapolated. In some instances, points from isotope dilution were negative or close-to-zero corrected values in which case they were excluded from the analysis. This amounted to max two data points per curve leaving at least three points in the saturated range. Protein determinations were made by the Bradford method.

### Photoaffinity labeling, affinity purification and mass spectrometry

#### Photoaffinity labeling (PAL) to hippocampal homogenate

To covalently label the native GHB high-affinity binding site, we developed an equilibrium PAL protocol in rat hippocampal homogenate (expressing up to 60 pmol/mg protein of target protein). A photoligand (**3**) was designed for this specific purpose, carrying two orthogonal azide moieties: one for photolinking and one for biotin tagging. Rat hippocampal homogenate was prepared as described for cortical tissue and incubated at a protein concentration of 0.125 mg/mL with 600 nM **3** for 60 min at 4 °C in the dark (concentration corresponds to 10x the IC_50_ value obtained in the [^3^H]-**5** radioligand binding assay). For the competition experiments that enable differentiation of endogenously biotinylated proteins from the biotin-labeled GHB target, **2** was added during this incubation step in increasing concentration. The homogenate was then transferred onto non-tissue culture-treated polystyrene plates and irradiated for 4 min at room temperature using a UVP Benchtop transilluminator (Thermo Fisher) set to high intensity (302 nm, 8 W, M-20 V) for photo-crosslinking to the aromatic azide group of **3**. Excess **3** was subsequently washed away using 1x PBS and centrifugation. For Staudinger-Bertozzi ligation to the alkylic azide group, the membranes were resuspended to a protein concentration of 0.5 mg/mL in 1x PBS and solubilized with 0.1% SDS and 1 mM EDTA for 15 min at 37 °C. EZ-Link™ Phosphine-PEG_3_-Biotin (#88901, Thermo Fisher) was added to a final concentration of 200 μM and the reaction was allowed to proceed for 60 min at 37 °C under shaking. Prior to streptavidin affinity enrichment, excess EZ-Link™ Phosphine-PEG3-Biotin was removed using PD MiniTrap G25 spin columns (#GE28-9180-07, GE Healthcare Biosciences, Pittsburgh, PA, USA).

#### Streptavidin affinity enrichment

Biotinylated proteins were enriched using Pierce™ High Capacity Streptavidin Agarose (#20361, Thermo Fisher). The solubilized membranes were diluted to a final concentration of 0.01% SDS and incubated with the resin under rotation for 30 min at room temperature. Enrichment was followed by a rigorous washing procedure (3 x 1 min with 10 column volumes (CV) PBS, 0.01% Tween-20 and 3 x 10 min with 10 CV PBS, 0.01% Tween-20). Biotinylated proteins were eluted by boiling in 1x NuPAGE™ LDS sample buffer (#NP0007, Thermo Fisher) supplemented with 100 μM DL-dithiothreitol (DTT) at 100 °C for 10 min under vigorous shaking. Eluates were loaded onto NuPAGE™ 4-12% Bis-Tris gels (Thermo Fisher) and run for 50 min at 175 V. Gels were stained with GelCode™ Blue Stain (#24590, Thermo Fisher) according to the manufacturer’s instructions. Gel sections between the 70 kDa and 25 kDa marker (PageRuler™ Prestained Protein Ladder, 10 to 180 kDa (#26616, Thermo Fisher) were cut out and diced into 1x1 mm cubes. After destaining using a 1:1 mixture of 5 mM ammonium bicarbonate pH 8.5 (ABC) and 100% acetonitrile for 10 min at room temperature, gel pieces were washed three times with ABC and 100% acetonitrile each. Proteins were reduced using 10 mM DTT for 10 min at 40 °C, followed by alkylation using 50 mM iodoacetamide (Sigma-Aldrich) for 20 min at room temperature in the dark. Samples were in-gel digested using 70 ng/band endoproteinase Lys-C (Sigma-Aldrich) overnight at 37 °C and 175 ng/band trypsin (Sigma-Aldrich) for 8 h at 37 °C. After acidification to a concentration of 1% formic acid, peptide extracts were loaded onto in-house packed C18 STAGE Tips (#14-386, 3M™ Empore) and eluted into a 96-well microtiter plate with 2 x 20 μL 40% acetonitrile, 0.5% acetic acid in water, followed by removal of organic solvents in a vacuum centrifuge and reconstitution of peptides in 2% acetonitrile, 0.5% acetic acid, 0.1% TFA in H_2_O.

#### Anti-biotin Western blotting

To verify the photoaffinity labeling of GHB high-affinity binding sites by **3** and competition by **2**, anti-biotin Western blotting using HRP-streptavidin was performed. Samples were prepared by adding 4x NuPAGE™ sample buffer (Thermo Fisher) and 100 mM DTT. Samples were denatured for 10 min at 40 °C, followed by centrifugation for 5 min at 11,000 x *g*. Samples were loaded onto Mini-PROTEAN^®^ TGX™ gels (Bio-Rad) and run for 40 min at 200V in 1x Tris/glycine/SDS (25 mM Tris, 192 mM glycine, 0.1% SDS, pH 8.3) running buffer. Proteins were transferred to a PVDF membrane using the Trans-Blot^®^ Turbo™ System (#1704156, Bio-Rad) and blocked with 3% BSA in 1x tris-buffered saline (TBS) with 0.1% Tween-20 detergent (TBS-T) for 1 h at room temperature. The membrane was then incubated with 0.5 mg/L streptavidin-HRP in 1% BSA in 1x TBS-T for 30 min at room temperature, followed by three quick washes with milliQ H_2_O and three washes with 1x TBS-T for 20 min at room temperature. The membrane was developed using Pierce ECL western blotting substrate (#32106, Thermo Fisher) for 4 min in the dark. Chemiluminescence was read using a FluorChem HD2 (Alpha Innotech, San Leandro, CA, USA).

#### LC-MS/MS

All samples were analyzed on an Easy-nLC 1000 coupled to a Q-Exactive HF instrument (Thermo Fisher) equipped with a nanoelectrospray source. Peptides were separated on a 15 cm analytical column (75 μm inner diameter) and packed in-house with 1.9 μm C18 beads (Dr. Maisch, Germany). The column temperature was maintained at 40 °C using an integrated column oven (PRSO-V1, Sonation GmbH, Biberach, Germany). Peptides were separated by a linear gradient of increasing acetonitrile in 0.5% acetic acid for 35 min with a flow rate of 250 nL/min. The Q-Exactive HF mass spectrometer was operated in data-dependent acquisition mode. Spray voltage was set to 2 kV, S-lens RF level at 50, and heated capillary temperature at 275 °C. All experiments were performed in the data-dependent acquisition mode to automatically isolate and fragment Top10 multiply-charged precursors according to their intensities. Former target ions were dynamically excluded for 40 sec, and all experiments were acquired using positive polarity mode. Full scan resolution was set to 60,000 at m/z 200 and the mass range was set to m/z 350-1400. Full scan ion target value was 3E6 allowing a maximum fill time of 100 ms. Higher-energy collisional dissociation (HCD) fragment scans were acquired with optimal setting for parallel acquisition using 1.3 m/z isolation width and normalized collision energy of 28. Ion target value for HCD fragment scans were set to 1E5 with a maximum fill time of 45 ms and analyzed with 60,000 resolution.

#### LC-MS/MS data analysis

Raw LC-MS/MS data was processed using the MaxQuant software (v. 1.5.5.1) and further data analysis was performed using Perseus (v. 1.5.6.0, Max-Planck Institute of Biochemistry, Department of Proteomics and Signal Transduction, Munich, Germany), Microsoft Office Excel and GraphPad Prism (v. 7). The MaxQuant search against the rat and mouse UniProt databases (downloaded 13.03.2017), leading to the identification of 1184 protein hits. In addition, the default contaminant protein database was included and any hits to this were excluded from further analysis. Further, proteins only identified by site were excluded. Carbamidomethylation of cysteine was specified as a fixed modification; phosphorylation of serine, threonine and tyrosine residues, oxidation of methionine, pyro-glutamate formation from glutamine and protein N-terminal acetylation were set as variable modifications. Proteins were quantified using the label free quantification (LFQ) algorithm ^42^. The LFQ intensity of proteins selectively labeled with **3** should decrease in a concentration-dependent manner with increasing concentrations of the competing ligand **2**. Unspecific binding or endogenously biotinylated proteins should not show this concentration-dependent behavior. Proteins only identified by site or from the reverse database were excluded. Data were then exported after initial filtering in Perseus to GraphPad Prism, and non-linear regression was performed for all proteins using the ‘*One site–Fit logIC_50_*’ function. Best-fit values for Top (upper plateau of the concentration-response curve) were plotted against the coefficient of determination (R^2^) values to identify proteins with competitive concentration-dependence behavior (*refer to Extended Data Table 1a*). Supplementary Table 1 contains entire data set.

### Validation of CaMK2α as the GHB high-affinity target

#### *In vitro* autoradiography in knockout (-/-) brain slices

*In vitro* autoradiography was performed on slices from adult male *Camk2a* -/- and *Camk2b* -/- mice. Wildtype (WT; +/+) littermates were used as controls. Initially, mice were euthanized by cervical dislocation, brains were dissected out quickly and snap frozen on liquid nitrogen. Brains were stored at -80 °C until further processing. On the day of the assay, brains were allowed to thaw for 30 min before they were cut on a CM1860 cryostat (Leica, Wetzlar, Germany) in 12 μm coronal sections at -20 °C. Sections were thaw-mounted on Super Frost Plus slides (Thermo Fisher). Before further processing, sections (3-4 per slide) were thawed for 1 h at room temperature. Subsequently, sections on slides were pre-incubated in binding buffer (50 mM KH_2_PO_4_ buffer, pH 6.0) for 30 min at room temperature under gentle shaking (20 rpm). Radioligand incubation followed for 1 h in binding buffer containing 1 nM [^3^H]-**1** or 7 nM [^3^H]-**5** at room temperature under gentle shaking (20 rpm). For [^3^H]GHB binding, sections were incubated for 30 min with a radioligand concentration of 30 nM at 4 °C. For determination of NSB, buffer/radioligand solution was supplemented with a non-radioactive displacer structurally unidentical to the radioligand employed (either 1 mM GHB or **1**). Reactions were terminated by washing 2 x 20 sec in ice-cold buffer followed by 3 dips in ice-cold dH_2_O. Sections were then air-dried for 1 h and placed in a fixator containing paraformaldehyde (PFA) vapour overnight at room temperature. The next day, sections were transferred to a desiccator box containing silica gel for 3 h. Finally, slides were exposed to a BAS-TR2040 Imaging Plate (Science Imaging Scandinavia AB, Nacka, Sweden) for 3 days in a radiation-shielded imaging plate cassette at room temperature. The imaging plate was scanned on a CR-35 Bio scanner (Dürr Medical, Bietigheim-Bissingen, Germany). Section anatomy was validated using cresyl violet staining. Experiments were carried out on individual mice (*n* = 3-4) in four technical replicates.

Staining of exemplary slices was performed with the Tissue-Tek Manual Slide Staining Set (Sakura Finetek, Brøndby, Denmark). First, lipids were dissolved by passing sections through an ascending series of ethanol dilutions for 1 min each (50% ethanol:50% dH_2_O > 70% ethanol:30% dH_2_O > 100% ethanol > 100% ethanol). Rehydration followed by immersion in decreasing ethanol concentrations ending with pure dH_2_O (1 min each). Sections were then stained in 1% cresyl violet solution for 10 min. Subsequently, sections were rinsed in 0.07% acetic acid for 4-8 sec, washed in dH_2_O for 1 min and dehydrated in increasing concentrations of ethanol for 30 sec each. Finally, sections were transferred to a xylene bath until mounting of coverslips with DPX mounting media. Slides were dried overnight, and images were acquired using a 1.25x objective under a bright-field microscope and sellSens software (Olympus Life Science, Ballerup, Denmark).

#### Radioligand binding assay on knockout (-/-) tissue crude homogenates (*Extended Data Fig. 2H*)

Crude homogenates of cortical tissues from adult male *Camk2a* -/-, *Camk2b* -/- and corresponding +/+ litter mates were individually prepared to generate biological replicates. In brief, brain tissues were quickly dissected after decapitation and stored in liquid nitrogen. Tissues were thawed in ice-cold binding buffer (50 mM KH_2_PO_4_ buffer, pH 6.0) (5x volume/weight) followed by homogenization using 2 x 1 mm zirconium beads in a Bullet Blender (NextAdvance, NY, USA) for 20 sec at max speed. Homogenates were centrifuged at 11,000 x *g* for 2 min at 4 °C, supernatant removed and the pellet resuspended in ice-cold binding buffer. Washing was repeated twice and protein concentration of homogenates was determined using the Bradford method, aliquoted and stored at -20 °C until further use. Radioligand binding assays were performed exactly as described for rat cortical homogenate using [^3^H]-**1** 5 nM.

#### PAL on knockout tissue homogenates (*Extended Data Fig. 2L-Q*)

To further verify the observed selectivity for CaMKIIα, **3** was tested using the same crude homogenates from *Camk2a* and *2b* -/- and +/+ tissues. The protocol was conducted exactly as described above for PAL to hippocampal homogenates followed by anti-biotin Western blotting.

#### Cell culturing and transfection of CaMKII in HEK293T cells

HEK293T cells were maintained in DMEM GlutaMAX medium (#61965026, Gibco) supplemented with 10% fetal bovine serum and 1% penicillin-streptomycin (#15140122, Invitrogen), in a humidified 5% CO_2_ atmosphere at 37 °C. Transfections with various CaMKII constructs were performed using PEI (Polysciences Inc., Warrington, PA, USA). The day before transfection, cells were seeded in 15 cm culture dishes at a density of 4.5 × 10^6^. On the day of transfection, 16 μg plasmid DNA was diluted in 2 mL serum-free medium and 48 μL 1 mg/mL PEI added. After 15 min of incubation at room temperature, the DNA/PEI mixture was added to the cells. Transfected cells were used for Western blotting or [^3^H]-**1** radioligand binding experiments.

#### Western blotting (*Extended Data Fig. 3*)

To check for successful transfection and expression of CaMKII constructs in HEK cells, Western blotting was performed as described for anti-biotin Western blotting. Membranes were probed with two primary antibodies targeting the myc-tag or CaMKIIβ (both diluted 1:1,000 and stored at -20 °C) and Na^+^/K^+^-ATPase (diluted 1:10,000 and stored at -20 °C) for 1 h at room temperature followed by 3 x 5 min washes with TBS-T. Subsequently, membranes incubated with species-specific secondary antibody, anti-mouse-HRP and anti-rabbit-HRP (1:2,000, 1 h at room temperature, stored at 4 °C) followed by 3 x 10 min washes with TBS-T. All antibodies were prepared in 1% (w/v) BSA in TBS-T and re-used up to four times. Membranes were probed with 1:1 mixture of ECL Detection Reagent (GE Healthcare) for up to 4 min before capturing the luminescence signal using a FluorChem HD2 (Alpha Innotech).

#### [^3^H]-**1** binding assay to HEK293T cell homogenates

Cell homogenates were prepared 48 h post-transfection by washing the cells with ice-cold PBS and harvesting by scraping. Cells were collected and centrifuged for 10 min at 1,500 x *g*. Cell pellets were resuspended in ice-cold binding buffer (50 mM KH_2_PO_4_ buffer, pH 6.0), and homogenized using 2 x 1 mm zirconium beads in a Bullet Blender for 20 sec at max speed. Protein concentration was determined using the Bradford method. Aliquots were stored at -20 °C until the day of assay.

Binding experiments were performed in a 48-well setup which was optimized to a protocol including 100-200 μg protein per well, 40 nM [^3^H]-**1** for competition binding and test compound in 400 μL total volume of binding buffer (50 mM KH_2_PO_4_ buffer, pH 6.0). Either WT or mutant CaMKII whole-cell homogenates were used for the competition binding. NSB was determined with 30 mM GHB. Saturation experiments employed [^3^H]-**1** in concentrations 10-10,000 nM (isotope dilution above 500 nM). Equilibrium binding was achieved by incubation for 60 min at 0-4 °C. To permit filtration, proteins were precipitated by addition of ice-cold acetone (4x of assay volume) followed by vortexing and incubation at -20 °C for 60 min. Protein-ligand bound complexes were collected by rapid filtration through GF/C unifilters (Whatman Schleicher and Schuell, Keene, NH) using a Brandell M48-T cell harvester (Alpha Biotech) and rapid washing with binding buffer. Radioactivity counts (DPM) was measured in a Packard Tricarb 2100 liquid scintillation counter using 3 mL of OptiFluor scintillation liquid (PerkinElmer) and counting for 3 min per sample.

#### Data analysis

Binding data were analyzed as described above for rat cerebral cortical homogenate. All experiments were replicated in at least three individual experiments with technical triplicates using at least three different batches of cell homogenate. IC_50_ values were converted to *K*_I_ values by means of the Cheng-Prusoff equation. *K*_D_ values were obtained from saturation experiments. 0-1 single data points of the individual saturation experiments were excluded from the analyses using same exclusion criteria as mentioned in the homogenate data analysis section.

### Localization of binding site to the CaMKIIα hub domain

#### Expression and purification of the CaMKIIα hub domain

The human CaMKIIα hub domain (UniprotKB Q9UQM7, residues 345-475), WT or containing six mutations (Thr354Asn, Glu355Gln, Thr412Asn, Ile414Met, Ile464His, and Phe467Met, referred to as the 6x Hub), was expressed and purified similar as to previously described ^38,43^. Briefly, the hub domain with an N-terminal 6His-prescission protease expression tag was inserted into a pSKB2 vector with kanamycin resistance. BL21 (DE3) *E. coli* cells transformed with the vector were cultured in TB media supplemented with phosphates. Protein expression was induced at OD_600_ = 0.6-0.8 by addition of 1 mM IPTG. Expression proceeded for 15-18 h while shaking at 18 °C. All subsequent purification steps were performed at 4 °C and all columns were made by GE Healthcare unless otherwise noted.

Induced cells were pelleted, resuspended in buffer A (25 mM Tris, 150 mM KCl, 50 mM imidazole, 0.5 mM DTT, 10% (v/v) glycerol, pH 8.5 at 4 °C), and lysed using a cell disrupter. Soluble cell lysate was passed over a 5 mL Nickel IMAC column. The column was washed with buffer A, and immobilized protein was eluted with a mixture of 25% buffer A and 75% buffer B (25 mM Tris, 150 mM KCl, 1 M imidazole, 10% glycerol, pH 8.5 at 4 °C). A HiPrep 26/10 column was used to exchange the protein into buffer C (25 mM Tris, 150 mM KCl, 10 mM imidazole, 1 mM DTT, 10% glycerol, pH 8.5 at 4 °C). Prescission protease was added overnight to remove the 6His tag. The protein was then concentrated using Amicon Ultra filters with a molecular weight cutoff of 50 kDa. A final purification step was performed using a Superose-6 gel filtration column equilibrated with the final protein storage buffer (25 mM Tris, 150 mM KCl, 10% (v/v) glycerol, 2 mM DTT, 1 mM TCEP, pH 8.0 at 4°C). Fraction purity was assessed by SDS-PAGE. Sufficiently pure fractions were pooled, concentrated, and flash-frozen in liquid nitrogen for storage at -80 °C.

#### Co-crystallization of compound **2** with the 6x Hub

Crystals were grown via sitting drop vapor diffusion at 20 °C. Reservoir solution contained 20% w/v PEG3350, 200 mM potassium acetate, pH 8.1. Protein stock concentration was 17 mg/mL. Compound **2** was dissolved in DMSO to 100 mM and 5 μL was added to 50 μL of protein solution for a final **2** concentration of approximately 10 mM, 10x the protein molarity. The protein and **2** were incubated at 4 °C for 2 h prior to dispensing. 200 nL drops were dispensed by adding 100 nL of protein stock to 100 nL of reservoir solution. Drops were equilibrated against 45 μL of reservoir solution. Crystals were harvested and frozen in cryoprotectant (20% w/v PEG3350, 20% v/v glycerol, 100 mM HEPES pH 7.0) 7 days after trays were set. X-ray diffraction data was collected at the Advanced Light Source, Beamline 8.3.1, at wavelength 1.115830 Å and temperature 100 K.

#### Structure determination

Data were processed using XDS^44^ and scaled and merged with Aimless^45^ in the CCP4 suite^46^. The space group was C2221 with cell dimensions a = 101.43, b = 182.96, c = 106.47, α = β = γ = 90°. Phases were determined by molecular replacement using Phenix Phaser^47^. A previously determined structure of the protein in an unliganded state (PDB entry 6OF8) was used as the search model^38^. Phenix Refine^47^ was used for structure refinement and Coot^48^ was used for model building. The cif dictionary file for **2** was generated using Phenix Elbow^49^. Software used in this project was curated by SBGrid^50^. Electron density in the shape of the compound was seen at the hypothesized binding sites of 6 of the 7 subunits in the crystallographic asymmetric unit prior to any structure refinement. Partial occupancy of the ligand at the binding site of the seventh subunit (chain A, the lone ‘Trp in’ subunit) was observed but not modeled (*Extended Data Fig. 5A-I*).

#### Tabulated x-ray crystallography data collection and refinement statisticsStatistics for the highest-resolution shell are shown in parentheses

Tabulated x-ray crystallography data collection and refinement statistics
|  | **6x Hub with compound 2** |
| --- | --- |
| **Wavelength** | 1.11583 |
| **Resolution range** | 106.47 - 2.20 (2.27 - 2.20) |
| **Space group** | C 2 2 21 |
| **Unit cell** |  |
| a, b, c | 101.43, 182.96, 106.47 |
| $\alpha$, $\beta$, $\gamma$ | 90, 90, 90 |
| **Total reflections** | 671271 (58251) |
| **Unique reflections** | 50585 (4349) |
| **Multiplicity** | 13.3 (13.4) |
| **Completeness (%)** | 100 (100) |
| **Mean I/sigma(I)** | 22.7 (0.8) |
| **Wilson B-factor** | 61.30 |
| **R-merge** | 0.072 (3.621) |
| **R-meas** | 0.079 (3.933) |
| **R-pim** | 0.021 (1.066) |
| **CC1/2** | 1 (0.292) |
| **Refinement Resolution** | 91.48 - 2.20 (2.28-2.20) |
| **Reflections used in refinement** | 50548 (4985) |
| **Reflections used for R-free** | 1715 (166) |
| **R-work** | 20.98 (36.35) |
| **R-free** | 24.37 (36.67) |
| **CC(work)** | 0.958 (0.568) |
| **CC(free)** | 0.957 (0.628) |
| **Number of non-hydrogen atoms** | 7511 |
| **macromolecules** | 7303 |
| **ligands** | 121 |
| **solvent** | 87 |
| **Protein residues** | 899 |
| **RMS(bonds)** | 0.003 |
| **RMS(angles)** | 0.56 |
| **Ramachandran favored (%)** | 96.91 |
| **Ramachandran allowed (%)** | 3.09 |
| **Ramachandran outliers (%)** | 0.00 |
| **Rotamer outliers (%)** | 2.69 |
| **Clashscore** | 3.88 |
| **Average B-factor** | 82.50 |
| **macromolecules** | 82.92 |
| **ligands** | 72.84 |
| **solvent** | 61.19 |
| **Number of TLS groups** | 37 |

Statistics for the highest-resolution shell are shown in parentheses.

### Biophysical and biochemical CaMKIIα assays

#### Surface plasmon resonance (SPR) biosensor analysis (*Extended Data Fig. 4A-E*)

SPR measurements were performed at 25 °C using a Pioneer FE instrument (Molecular Devices, FortéBio). 6x Hub human protein, WT hub or full-length human recombinant CaMKIIα (#02-109, Carna Biosciences) were immobilized on to a biosensor surfaces by amine coupling using a 20 mM NaAc pH 5 immobilization buffer. HBS-P (10 mM Hepes, 150 mM NaCl, 0.005% Tween-20, 1 mM DTT) pH 7.4 or pH 6 running buffer were used for all experiments with CaMKIIα hub, while for experiments with full-length CaMKIIα, HBS-P buffer pH 7.4 supplemented with CaCl_2_ (Ca^2+^) (500 μM) was used. The ligands were injected in 2-fold serial dilution over sensor chip surface with immobilized proteins. Control peptide **7** and calmodulin (CaM) were used as positive controls to evaluate activity of the SPR assay with CaMKIIα hub and full-length protein, respectively. Between CaM injections, the biosensor chip surface was regenerated by injections of HBS-P buffer supplemented with 100 μM EDTA. The data were analyzed using Qdat Data Analysis Tool version 2.6.3.0 (Molecular Devices, FortéBio). The sensorgrams were corrected for buffer bulk effects and unspecific binding of the samples to the chip matrix by blank and reference surface subtraction (flow cell channel activated by injection of EDC/NHS and inactivated by injection of ethanolamine). The dissociation constants (*K*_D_) were estimated by plotting responses at equilibrium (R_eq_) against the injected concentration and curve fitted to a Langmuir (1:1) binding isotherm.

#### Differential scanning fluorimetry (DSF) (*Extended Data Fig. 4F-H*)

Thermal melting points (T_m_) of the CaMKIIα WT hub with and without the presence of GHB, **1** or **2** were assessed by DSF measured on a Mx3005P qPCR System (Agilent Technologies, Waldbronn, Germany). Samples were prepared in qPCR 96-well plates (25 μL/well) with a final concentration of 0.1 mg/mL CaMKIIα and 8x SYPRO^®^ Orange Protein Gel Stain (#S6650; Life Technologies) in MES buffer (20 mM MES, 150 mM NaCl, 1 mM DTT; pH 6). GHB, **1** and **2** were tested in 3-fold dilution series. Fluorescence was monitored using excitation at 492 nm and emission at 610 nm from 25–114 °C in 90 cycles with a 1 °C temperature increase per min. Data was processed in GraphPad Prism (v. 8). T_m_ values were calculated by fitting the sigmoidal curves of normalized fluorescence intensity versus temperature to the Boltzmann equation. The difference in T_m_ (ΔT_m_) of each compound concentration compared to CaMKIIα WT hub was plotted as concentration-response curves and maximum ΔT_m_ was derived from nonlinear regression using the one site model by non-linear regression. Experiments were performed in at least three independent experiments using singlicates.

#### Computational modeling (*Extended Data Fig. 5J*)

All computational work was performed using the Maestro Schrödinger package (Schrödinger Release: 2018-1). The co-crystal structure complex of CaMKIIα-compound **2** was prepared using the Protein Preparation Wizard^51^ including hydrogen optimization at pH 6.4 of the ionizable polar groups using PROPKA^52^. The PDB entry 5IG3^36^ was used to illustrate the ‘Trp-in’ observation. The Trp403 movement comparison was observed via a Protein Superimposition of the backbone atoms of both our reported ligand bound structure reported herein and that of the PDB entry 5IG3 (RMSD = 1.091 Å).

#### Intrinsic tryptophan fluorescence (Trp flip) assay (*Extended Data Fig. 5K-L*)

Human hub recombinant purified proteins and compounds (**1** and **2**) were diluted in assay buffer (10 mM HEPES pH 7.4, 150 mM NaCl, 1 mM DTT) and mixed in a microplate to obtain a protein concentration of 5 μM 6x Hub and 3.4 μM WT hub. For absorbance and background fluorescence measurements, compounds were mixed with buffer for each compound concentration. **1** and **2** showed no fluorescent properties. All measurements were performed in black half-area 96-well format low-binding OptiPlates (#6052260, PerkinElmer) for fluorescence and half-area UV-Star microplates (#675801, Greiner Bio-One) for absorbance. All measurements were recorded at 25 °C on a Safire^2^ plate reader (Tecan). Emission was recorded in the wavelength range of 300-450 nm with 1 nm increments and an excitation wavelength of 290 nm with 5 nm band widths. Fluorescence intensities at 340 nm were used for data analysis. To check for inner filter correction, the absorbance was measured in the range of 270-400 nm. **1** and **2** showed low absorbance and no inner filter effect.

The fluorescence intensities were normalized according to:

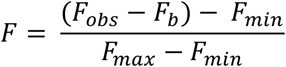

F_obs_ is the observed fluorescence intensity and F_b_ is the background fluorescence for compound in buffer alone. F_max_ is the fluorescence intensity of hub alone without compound, and F_min_ is the fluorescence intensity when plateau is reached at high compound concentrations in the presence of hub. Since **1** did not reach a plateau at high compound concentrations, F_min_ was set to the fluorescence intensity of buffer for all compounds tested. Fluorescence intensities usually spanned from 10,000-55,000 for **2** and 36,000-59,000 for **1**, while the fluorescence intensity for buffer was around 1000. Non-linear regression was used for curve-fitting using the equation for ‘*log(inhibitor) vs. response with variable slope*’ to determine IC_50_ values (GraphPad Prism, v. 8). All curves are pooled data (means ± SEM) each performed in technical triplicates using at least two different batches and several different aliquots of protein. For 6x Hub (*n* = 8) and for WT hub (*n* = 2-5).

#### ADP-Glo Assay (*Extended Data Fig. 7E*)

Syntide-II substrate phosphorylation was assayed using the ADP-Glo Assay kit (#V9101, Promega) using CaMKIIα full-length protein with a N-terminal GST-tag (#02-109, Carna Biosciences) or CaMKIIα full-length protein with a C-terminal 6xHis-tag (#PR4586C, Thermo Fisher). All experiments were performed in 384-well white polypropylene plates (#784075, Greiner) with a working volume of 25 μL. Kinase detection reagent was prepared according to manufacturer’s protocol. Linearity of the assay was assessed using various concentrations of ADP to ATP mixture in the ADP-Glo kinase reaction buffer (40 mM Tris (pH 7.5), 0.5 mM CaCl_2_, 20 mM MgCl2, 0.1 mg/mL BSA, 30 μg/mL calmodulin (#P1431, Sigma-Aldrich), 50 μM DTT) as described in the manufacturer’s protocol. For inhibitor studies, the total assay volume was 20 μL, the volume of the kinase reaction was 5 μL (1 μL 5x inhibitor + 2 μL 2.5x CaMKIIα (= 3 ng final) + 2 μL 2.5x substrate mix (= 25 μM ATP and 50 μM syntide-II final concentrations). Kinase reactions were incubated 60 min at 37 °C, after this 5 μL ADP-Glo reagent was added to each well and incubated for 40 min at room temperature. Finally, 10 μL kinase detection reagent was added to each well and after 30 min of incubation at room temperature, luminescence was recorded on an EnSpire^®^ Multimode Plate Reader (PerkinElmer). Kinase reactions without enzyme or substrate were used as controls. Curve fitting and data analysis was done using GraphPad (v. 8), and IC_50_ values were determined using the equation for *log(inhibitor) vs. response with variable slope*. All curves are pooled data (means ± SEM) of three independent experiments each performed in technical triplicates.

### *In vitro* assays using cultured neurons and slices

#### Primary neuronal cultures

Cortical neuronal cultures were prepared from E16-E18 embryos. Tissue dissociation and neuronal isolation (approximately 6-9 embryos per female mouse) was performed using validated antibodies with MACS^®^ technology according to the manufacturer’s description (#130-094-802 and #130-115-389, Miltenyi Biotec). When more than 6×10^7^ cells were obtained from tissue dissociation, two separation columns were used in neuronal isolation. As a result, approximately 2×10^7^ – 3×10^7^ cells were frequently obtained. Primary hippocampal neuronal cultures were prepared according to the procedure described in Goslin and Banker (1991) ^53^. Briefly, hippocampi were isolated from brains of E16.5 embryos and collected altogether in 10 ml of Neurobasal™-A medium (NB) (#10888-022, Invitrogen) on ice. After two washings with NB, the neurons were dissociated with Gibco™ trypsin/EDTA solution (#25300054, Thermo Fisher). Isolated cortical and dissociated hippocampal neurons were resuspended in NB supplemented with 2% B-27™ Supplement (#17504044, Invitrogen), 1% GlutaMAX™ Supplement (#35050038, Invitrogen) and 1% penicillin-streptomycin (Invitrogen), and plated in poly-D-lysine-coated 96-well, 12-well or 24-well dishes (~60,000, 100,000 or 400,000 cells/well). Cultures were maintained at 37 °C and 5% CO_2_ and fed every 48-62 h, where half of the conditioned media were replaced with fresh media. To optimize culturing time (cortical cultures) sufficient CaMKIIα expression, mRNA and protein levels were assessed by qPCR and Western blot, respectively.

#### Validation of mRNA levels in cortical neuronal cultures (*Extended Data Fig. 6A*)

To determine the expression level of *Camk2a* and *Camk2b* during culturing time days-in-vitro (DIV) 4-14, mRNA levels were verified by reverse transcription quantitative polymerase chain reaction (RT-qPCR). Cortical neuronal cultures plated in 24-well dishes were harvested in RLT plus Lysis buffer (Qiagen) and stored at -80 °C prior to RNA extraction. Each sample was homogenized with zirconium oxide beads (2 x 2 mm) using a Bullet Blender and total RNA was extracted with RNeasy Plus Mini kit (#74136, Qiagen) and PureLink^®^ DNase (#12185010, Invitrogen) according to the manufacturer’s instructions with minor changes. Total RNA amounts were quantified using NanoDrop 2000 (Thermo Fisher). 500 ng of RNA was reverse-transcribed using qScript^®^ cDNA Supermix (Quanta Bio) on an Eppendorf Mastercycler Personal PCR machine. The cDNA was diluted to approximately 100 ng/μL and stored at -20 °C until RT-qPCR was performed. The PCR samples were prepared on ice by combining 5 μL cDNA, 10 μL Power SYBR green Master Mix (2x) (Applied Biosystems) and 0.15 μL of forward and reverse primers (*Camk2a* (F) 5’-GCTCTTCGAGGAATTGGGCAA-3’ (R) 5’CCTCTGAGATGCTGTCATGTAGT-3’, *Camk2b* (F) 5’GCACACCAGGCTACCTGTC-3’ (R) 5’-GGACGGGAAGTCATAGGCA-3’, *SDHA* (housekeeping gene) (F) 5’-GGAACACTCCAAAAACAGACCT-3’ (R) 5’-CCACCACTGGGTATTGAGTAGAA-3’; TAG Copenhagen) with distilled water to a final volume of 20 μL. Further, the RT-qPCR was initiated by 10 min heating to 95 °C followed by 40 PCR cycles of 15 s at 95 °C and 60 s at 60 °C on a Stratagene Mx3005P (Agilent Technologies). The cycle threshold (Ct) of the fluorescent PCR product was determined by default in MxPro qPCR Software (Agilent Technologies), and used to calculate the relative fold gene expression:

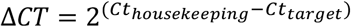

Data are given as means ± SEM from three independent cultures (*n* = 3) each performed in technical triplicates.

#### Validation of CaMKIIα protein levels by Western blotting (*Extended Data Fig. 6C-H*)

Protein expression levels (DIV 4-20) were examined by Western blot. Neurons (400,000 cells/well in 24-well dish) were harvested in 150 μL ice-cold radioimmunoprecipitation (RIPA) buffer supplied with a 1% protease/phosphatase inhibitor cocktail. Protein concentrations were determined with the Pierce BCA method, and neuronal samples were stored at -80 °C until Western blot analysis. Western blot samples were prepared by adding 4x Fluorescent Compatible Sample Buffer (Thermo Fisher), 100 mM DTT and harvested neuron samples to a final concentration of 1 μg/μL. Samples were heated for 10 min at 37 °C, followed by sonication and centrifugation for 2 min at 4 °C and 11,000 x *g*. 10 μg sample were loaded on 4-20% Mini-PROTEAN^®^ TGX™ gels (Bio-Rad) with iBright™ Prestained Protein Ladder (#LC5615, Invitrogen). SDS-PAGE was performed at 200 V for 40 min with Tris/glycine/SDS running buffer. Proteins were transferred to a low-fluorescence polyvinylidene difluoride (PVDF) membrane (Thermo Fisher) using the Trans-Blot^®^ Turbo™ transfer system (Bio-Rad). Membranes were blocked for 30 min at room temperature under constant agitation with 1x Blocker™ FL Fluorescent Blocking Buffer (#37565, Thermo Fisher). Membranes were probed with primary antibody targeting CaMKIIα (1:1,000, incubation overnight at 4 °C, stored at -20 °C), CaMKIIα-pThr286 (1:1,000, incubation overnight at 4 °C, stored at -20 °C) or GAPDH (1:10,000, incubation 1 h at room temperature, stored at 4 °C) followed by 3 x 5 min washes with TBS-T. Subsequently, membranes were incubated with species-specific secondary antibody (1:2,000, 1 h at room temperature, stored at 4 °C) followed by 3 x 10 min washes with TBS-T. All antibodies were prepared in 1% (w/v) BSA in TBS-T and re-used up to five times. Images were detected with the iBright FL1500 imaging system (Invitrogen) and quantified in Image Studio (Lite version 5.2). CaMKIIα-pThr286 signals were normalized to total CaMKIIα. Data are from one culture carried using three different wells (means ± SD).

#### Validation of CaMKIIα expression by [^3^H]-**1** radioligand binding (*Extended Data Fig. 6B*)

For [^3^H]-**1** radioligand binding, [^3^H]-**1** (40 nM) and GHB (8 mM) for NSB. Cells grown in 6-cm dishes (DIV 10) were harvested in ice-cold binding buffer (50 mM K_2_PO_4_, pH 6.0) and a crude homogenate prepared. Briefly, cells were collected in Eppendorf tubes, subjected to three centrifugation (11,000 x *g*)-resuspension steps and Bullet Blender homogenization. Membranes were diluted to give 10-20 μg final protein per well. Data are from three different cultures carried out in technical replicates (means ± SEM).

#### pThr286 Autophosphorylation in cultured cortical neurons

Cortical neuronal cultures (DIV 18-20) plated in 24-well dishes were incubated for 1 h (37 °C, 5% CO_2_) with the desired compounds diluted in assay buffer (HBSS supplied with 20 mM HEPES, pH 7.4). Buffer was used for measurement of basal and Ca^2+^ as stimulation control, typically 5 mM. Subsequently, neurons were harvested in 150 μL ice-cold RIPA buffer supplied with a 1% protease/phosphatase inhibitor cocktail. Protein concentrations were determined using Pierce BCA method, and neuronal samples were stored at -80 °C until Western blot analysis targeting CaMKIIα and CaMKIIα-pThr286 as described in the section ‘Validation of CaMKIIα protein expression’. Statistical analysis was performed with GraphPad Prism (v. 8). If groups had equal variances, One-Way ANOVA was used, followed by Dunnett’s post-hoc test comparing Ca^2+^ stimulation with treatments. If no equal variance was obtained, statistical analysis was done with Brown-Forsythe and Welch ANOVA, followed by Dunnett’s T3 multiple comparisons test to compare basal and treatment. Exact number of replications/batches of cultures are stated in the figure legends.

#### Excitotoxic stimulation

Cortical neuronal cultures (DIV 16-18) were exposed to a pathological stimulus of 1-200 μM L-glutamate (Glu) plus 20 μM glyine (Gly) for 1 h (37 °C, 5% CO_2_). HBSS supplemented with HEPES (pH 7.4) was used as assay buffer. The neuronal media was replaced with 50 μL assay buffer and an equal volume of assay buffer supplemented with 2x Glu/Gly was added. Subsequently, the cultures were washed with assay buffer and replaced with fresh neurobasal medium. When compounds were added during the excitotoxic insult, half of the assay buffer was replaced with buffer containing 1x Glu/Gly and 2x compound. With application of compounds following excitotoxic insult, half of the fresh neurobasal medium was replaced with neurobasal medium containing 2x compound. All compound incubations were 1 h at 37 °C and 5% CO_2_. **1** was used in the indicated concentrations. Tat-**4** (10 μM) was used as a positive control. Finally, the neurons were washed with assay buffer and incubated 20-24 h at 37 °C and 5% CO_2_ in NB until further assay analysis.

#### Cell viability lactate dehydrogenase (LDH) assay

Following 20-24 h Glu/Gly stimulation, neuronal cell death was assessed using the CyQuant™ lactate dehydrogenase (LDH) Cytotoxic Assay kit (#C20301, Invitrogen) according to the manufacturer’s instructions. Initially, lysis buffer (x10) was added in triplicate to determine the maximum LDH release. Following 45 min incubation at 37 °C, 5% CO_2_, 50 μL from each well was transferred to a 96-well flat-bottom ELISA plate (#655101, Greiner). Triplicates or quadruplicates were performed with each condition. To initiate the coupled enzymatic reactions, 50 μL reaction mixture was added to each sample well. After 30 min incubation at room temperature and under protection from light, the enzymatic reactions were terminated with 50 μL stop solution. The background absorbance at 680 nm was subtracted from the LDH activity absorbance at 490 nm. Further, control wells containing only media were subtracted before further calculation of percentage LDH release compared to maximum cell death. One-Way ANOVA was used for statistical analysis in GraphPad Prism (v. 8), followed by Dunnett’s post-hoc test comparing control (Glu/Gly alone) with treatment. Exact number of replications/batches of cultures used are stated in the figure legends.

#### Co-localization studies in hippocampal neurons

Primary hippocampal neurons (DIV 14-19) plated in 12-well dishes were used for immunocytochemistry. All medium was removed and replaced with Tyrode’s solution (5 mM) with tetrodotoxin (TTX, 1 μM, #1078), NBQX (50 μM, #0373, both from Tocris Biosciences, Bristol, UK), and L-AP5 (10 μM, #A5282, Sigma-Aldrich) to block the basal activity of the neurons. After 10 min in H_2_O, tat-**4** (10 μM) according to ^54^, or compound **1** (2 mM) were added to the cultures for 20 min, followed by 2 min stimulation with Glu (400 μM). The neurons were fixated with 4% PFA/4% sucrose for 10 min at room temperature and washed 3 times 5 minutes in PBS. The coverslips with the fixated neurons were incubated in GDB buffer (0.2% BSA, 0.8 M NaCl, 0.5% Triton X-100, 30 mM phosphate buffer (PB), pH 7.4) with the primary antibodies: rabbit polyclonal GluN2B (1:100), mouse α-CaMKII (1:100) and guinea pig MAP2 (1:500) overnight in the dark at 4 °C. After incubation, coverslips were washed 3 times in PBS at room temperature, followed by a 1 h incubation at room temperature in GDB buffer with the secondary antibodies donkey anti-rabbit Alexa488 (1:100), donkey anti-mouse Cy3 (1:100), donkey anti-guinea pig Alexa647 (1:100). Finally, the coverslips were washed 3 times 5 min in DPBS, rinsed briefly in MilliQ, mounted with Mowiol^®^ 4-88 (#81381, Sigma-Aldrich) and dried overnight at 4 °C. An LSM700 Zeiss confocal microscope was used to make airyscan images of the neurons. Five to seven images were taken per coverslip. Colocalization of GluN2B and CaMKIIα in the secondary dendrites was determined using the Confined Displacement Algorithm (CDA) plugin in Fiji ImageJ (National Institutes of Health, USA). Data was normalized to the control group. For statistical analysis, GraphPad Prism (v. 8) was used. To show a significant effect of the treatments, One-Way ANOVA was used, followed by Dunnett’s post-hoc test comparing control with treatment. Exact number of replications/batches of cultures used are stated in the figure legends.

#### *Ex vivo* LTP recordings in mice (*Extended Data Fig. A-D*)

Adult male mice (> 8 w old) were used. After anesthesia with isoflurane (Nicholas Piramal) and decapitation, the brain was taken out quickly and submerged in ice-cold oxygenated (95%) and carbonated (5%) artificial CSF (ACSF; <4.0°) containing the following (in mM): 120 NaCl, 3.5 KCl, 2.5 CaCl_2_, 1.3 MgSO_4_, 1.25 NaH_2_PO_4_, 26 NaHCO_3_, and 10 D-glucose. Using a vibratome, 400-μm-thick sagittal slices were made. Hippocampal sections were dissected out afterward and maintained at room temperature for at least 1.5 h in an oxygenated and carbonated bath to recover before experiments were initiated. At the onset of experiments hippocampal slices were placed in a submerged recording chamber and perfused continuously at a rate of 2 mL/min with ACSF equilibrated with 95% O_2_, 5% CO_2_ at 30 °C. Extracellular recording of field EPSPs (fEPSPs) and stimulation were done using bipolar platinum (Pt)/iridium (Ir) electrodes (Frederick Haer). Stimulus duration of 100 μs for all experiments was used. In CA3–CA1 measurements, the stimulating electrode and recording electrode were placed on the CA3–CA1 Schaffer collateral afferents and apical dendrites of CA1 pyramidal cells (both 150–200 μm from stratum pyramidale), respectively. Upon placement of the electrodes, slices were given 20–30 min to rest before continuing measurements. Compound **1** (100 μM) was washed in 20 min before and continued to be washed in during the experiment. All paired-pulse facilitation (PPF) experiments were stimulated at one-third of slice maximum. Varying intervals were used in PPF: 10, 25, 50, 100, 200, and 400 ms. CaMKII-dependent LTP was evoked using a 100 Hz induction protocol (1 train of 1 s at 100 Hz, stimulated at one-third of slice maximum). During LTP slices were stimulated once per minute. Potentiation was measured as the normalized increase of the mean fEPSP slope for the duration of the baseline. Only stable recordings were included and this judgment was made blinded. Average LTP was defined as the mean last 10 min of the normalized fEPSP slope. Number of mice per experiment was at least *n* = 20 as specified in the figure legend and are given as means±SEM. For statistical analysis Repeated Measures ANOVA was used.

### *In vivo* and *ex vivo* studies

#### Study design, power analysis and statistics

All *in vivo* stroke studies were approved by the University of Otago Animal Ethics Committee and are reported according to the ARRIVE (Animal Research: Reporting In Vivo Experiments) guidelines. For behavioral experiments, six animals per group are required to achieve >80% power (86% calculated), considering the following parameters: α=0.05; with an effect size=1.5. For histological and immunohistochemical experiments, 5 animals per group are required to achieve >80% power (91% calculated), considering the following parameters: α=0.05; effect size 1; 3 concentrations; 2 groups, and correlation between measures =0.5. Parameters were determined from our prior work, in which we have demonstrated significant behavioural effects^55^, and on the assumption that variance was about 25%. It should be noted that more conservative effect sizes were used for these experiments, as it is harder to assess recovery over time between groups than looking at the effects of drug treatments on stroke size. Mice were allocated randomly to treatment groups, and the experiments performed blinded.

All statistical analyses relating to this section were performed using GraphPad Prism (v. 8). For *in vivo* studies, data are displayed as mean ± SD and plotted as box and whisker graphs, unless otherwise noted. For histological, electrophysiological and behavioral assessments poststroke, one-way and two-way ANOVA followed by Dunnett’s, Tukey’s or Bonferroni’s post-hoc test as well as two-tailed Student’s *t*-test were used when as appropriate and specified accordingly in the figure legends. For histological assessment of groups with an *n* = 5/group, one-way ANOVA followed by Kruskal-Wallis test was performed. P<0.05 was considered statistically significant.

#### Photothrombotic stroke

All procedures were performed in accordance with the guidelines on the care and use of laboratory animals set out by the University of Otago, Animal Research Committee and the Guide for Care and Use of Laboratory Animals (NIH Publication No. 85–23, 1996). All mice for the stroke studies were obtained from the Biomedical Research Facility, University of Otago, New Zealand.

Focal stroke was induced by photothrombosis in adult male C57BL/6*J* mice (3-4 months, 27-30 g) or aged female C57BL/6*J* mice (20-24 months, 32-40 g) as previously described^56^. Under isoflurane anesthesia (4% induction, 2-2.5% maintenance in O_2_) mice were placed in a stereotactic frame (9000RR-B-U, KOPF; CA, USA), and buprenorphine hydrochloride (0.1 mL of a 0.5 mg/kg solution, Temgesic^®^) was administered subcutaneously as pre-emptive post-surgical pain relief. Following sterilization of the skin using chlorhexidine (30% in 70% ethanol, Hibitane^®^), the skull was exposed through a midline incision, cleared of connective tissue and dried. A cold light source (KL1500 LCD, Zeiss, Auckland, New Zealand) attached to a 40x objective providing a 2-mm diameter illumination was positioned 1.5 mm lateral from bregma. Then, 0.2 mL of Rose Bengal (Sigma-Aldrich, Auckland, New Zealand; 10 mg/mL in normal saline) was administered i.p. After 5 min, the brain was illuminated through the exposed intact skull for 15 minutes, while keeping body temperature at 37 °C using a heating pad. The color temperature intensity used for all experiments to induce a stroke was 3300 K. The skin was glued and animals left in a cage placed on a heating pad during the wake-up phase. Sham surgery was performed in the exact same way, except that saline was injected instead of Rose Bengal. Mice were housed in groups of two to five under standard conditions in individually ventilated cages (IVC: Tecniplast): 21 ± 2 °C and humidity of 50 ± 10%, on a reverse 12 h light/dark cycle (white lights off from 07:00-19:00) with ad libitum access to food and water. Further, the mice were monitored and weighed on a daily basis. All animals were randomly assigned to a treatment groups and all assessments were carried out by observers blinded to the treatment group.

The sodium salts of GHB and **1** were dissolved in sterile dH_2_O. Drugs were administered i.p. as 10 mg/mL and 10 μL of solution per gram mouse body weight. Injection of compound (i.p.) was performed 30 min, 3 h, 6 h or 12 h after induction of the photothrombotic stroke. The vehicle groups received a corresponding volume of saline (0.9%).

#### Histological assessment

For quantification of the infarcted area, animals were deeply anaesthetized (i.p. pentobarbital 100 mg/kg) and transcardially perfused with saline followed by 4% PFA at either 3- or 7-days post-stroke. Brains were dissected out and post-fixed in 4% PFA for post-fixing overnight prior to being transferred into 30% sucrose solution for cryopreservation and kept at 4° C until sectioning. Brains were cut on a sliding microtome attached with a freezing stage (-21°C, Leica, Model 1300). A layer of sucrose solution (30%) was used to mount the brain onto the freezing stage and brains left to freeze (15-20 min) before 40 μm coronal sections were cut and collected into 24-well plates containing a cryoprotectant solution (0.1 M PBS, 30% sucrose, 1% polyvinylpyrrolidone, 30% ethylene glycol). Tissue was stored at -20 °C until required.

Every sixth section was washed in 0.05 M TBS solution for three consecutive 10 min washes. Sections were mounted on gelatin subbed slides, dried overnight, stained using 0.1% cresyl violet solution, passed through ascending concentration of alcohols, cleared in xylenes, and coverslipped using DPX mounting solution. Images of cresyl violet staining were then taken using an inverted montaging microscope (Model Ti2E Wideview, Nikon, Japan) set with a 2.5x objective lens. Images were then exported as TIFF files and opened on Fiji ImageJ to quantify infarct volume. Stroke volume was calculated as per the equation:

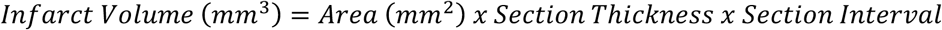

#### Behavioral assessment

Motor performance was determined using both the cylinder and grid-walking tasks as previously described^56^. All animals were tested one week prior to surgery on both behavioral tasks to establish baseline performance levels, and again seven days post-stroke, at approximately the same time each day, at the end of their dark cycle. All behaviors were scored by observers who were blind to the treatment group of the animals in the study as previously described^55^.

#### Grid-walking test

The grid-walking apparatus was manufactured using 12 mm square wire mesh with a grid area 32 cm / 20 cm / 50 cm (length / width / height). A mirror was placed beneath the apparatus to allow video footage in order to assess the animals’ stepping errors (i.e. ‘foot faults’). Each mouse was placed individually atop of the elevated wire grid and allowed to freely walk for a period of 5 min (measured in real time by stopwatch and confirmed afterwards by reviewing videotape footage). During this 5 min period, the total number of footfaults for each limb along with the total number of non-foot fault steps was counted and a ratio between footfaults and total steps taken calculated. Percent foot faults were calculated by: [#foot faults / (#foot faults + #non-foot fault steps) * 100]. To take into account differences in the degree of locomotion between animals and trials, a ratio between foot faults and total steps taken was used.

#### Cylinder task

The spontaneous forelimb task encourages the use of forelimbs for vertical wall exploration / press in a cylinder. When placed in a cylinder, the animal rears to a standing position, whilst supporting its weight with either one or both of its forelimbs on the side of the cylinder wall. A cylinder 15 cm in height with a diameter of 10 cm is used. Videotape footage of animals in the cylinder was evaluated quantitatively in order to determine forelimb preference during vertical exploratory movements. While the video footage was played in slow motion (1/5th real time speed), the time (sec) during each rear that each animal spent on either the right forelimb, the left forelimb, or on both forelimbs were calculated. Only rears in which both forelimbs could be clearly seen were timed. From these three measures, the total amount of time spent on either limb independently as well as the time the animal spent rearing using both limbs was derived. The percentage of time spent on each limb was calculated and these data were used to derive a spontaneous forelimb asymmetry index (% ipsilateral use / % contralateral use). The ‘contact time’ method of examining the behavior was chosen over the ‘contact placement’ method, as it takes into account the slips that often occur during a bilateral wall press post-stroke.

#### pThr286 autophosphorylation after photothrombotic stroke

Photothrombotic strokes were induced as described and brains collected. Mice were euthanized by cervical dislocation, followed by rapid extraction of the brain at 3, 6 and 12 h after stroke induction. Brains were snap-frozen and stored at -80 °C until further processing. On the day of the assay, brains were cut using a CM1860 cryostat (Leica) at -20 °C and the peri-infarct was collected using a tissue punch from the top quadrant of the left hemisphere. Tissue homogenization was achieved using a Bullet Blender in RIPA buffer supplemented with a 1% protease/phosphatase inhibitor cocktail. Protein concentrations were determined with the Pierce method. Samples were prepared for Western blot analysis by addition of 4x Fluorescent Compatible Sample Buffer (#LC2570, Thermo Fisher) and 100 mM DTT with a protein concentration of 2 μg/μL. Samples were heated for 5 min at 95 °C, sonicated and centrifuged 11,000 x *g* for 2 min at 4 °C. Western blotting and antibody incubation was performed exactly as described under ‘Validation of CaMKIIα protein’, however with 20 μg sample loaded. Experiments were carried out in 4-5 biological replicates using three technical replications. For statistics, One-way ANOVA, post-hoc Dunnett’s test was used.

#### Compound action potential (CAP) electrophysiological recordings from corpus callosum

Functional changes within the corpus callosum were measured by assessing compound action potential recordings (CAPs) based on Crawford *et al*^57^. Briefly, 14-days post-sham or stroke surgery, mice were anaesthetized with isoflurane, decapitated, and the brain rapidly removed. Coronal slices (400 μm thick) were cut with a vibratome (Campden Instruments model #MA752) in ice-cold artificial cerebrospinal fluid (aCSF (in mM): NaCl 124, KCl 5, NaH_2_PO_4_ 1.25, NaHCO_3_ 26, MgSO_4_ 1.3, CaCl_2_ 2, glucose 10; pH 7.4; gassed with Carbogen: 95% O_2_/5% CO_2_). Slices were then transferred to a holding chamber containing oxygenated aCSF at room temperature and were allowed to equilibrate under these conditions for at least 1 h prior to recording.

Brain slices were placed between nylon mesh nets in a commercial brain slice recording chamber (Kerr Scientific Instruments Ltd., Christchurch, NZ) that was maintained at room temperature, and constantly superfused at 1 mL/min with oxygenated aCSF. The slices were left undisturbed for 1 h before commencement of recordings. Electrical stimulation of the corpus callosum was performed using a bipolar microstimulating electrodes (50 μm diameter Teflon-coated tungsten wire), positioned over the corpus callosum in order to evoke CAPs. Biphasic electrical stimulation was applied as 0.1 msec pulses delivered at 0.1 Hz, at intensities ranging from 2 to 30 V. The evoked CAPs were recorded extracellularly at an approximate distance of 0.8-1.2 mm from the stimulation site using a crimped copper electrode inserted inside a glass microelectrode filled with aCSF. The recording electrode was attached to a KSI Bio Amplifier, with waveforms were filtered and amplified using a Powerlab 2/25 analog-digital converter (ADInstruments Pty., Sydney, Australia) and stored on a Macintosh computer hard drive for offline analysis using LabChart 7 Pro data acquisition software (ADInstruments Pty.). Waveforms were analyzed to assess peak responses for N1 (myelinated axons) and N2 (unmyelinated axons) and expressed as CAP responses (mV).

#### Biotinylated dextran amine (BDA) injection

Injection of BDA was carried out as we have previously described ^55^. On day 7 post-stroke, animals were anaesthetized and positioned in a stereotaxic frame (KOPF), and buprenorphine hydrochloride (0.1 mL of a 0.5 mg/kg solution, Temgesic^®^) was administered subcutaneously as pre-emptive post-surgical pain relief. Following sterilization of the skin using chlorhexidine (30% in 70% ethanol, Hibitane^®^), an incision was made to expose the skull and connective tissue cleared and the skull dried. A glass Hamilton syringe (5 mL; Hamilton Company, NV, USA) containing 10% BDA (10,000 MW; Invitrogen) was positioned above bregma (1.5 mm AP, 1.75 mm ML) to work out where to drill. Using a small drill (Micro 8v Max, Dremel), a 1 mm diameter burr-hole was carefully made through the skull. The Hamilton syringe was then lined back up over the burr-hole and lowered slowly through the skull into the left premotor cortex (0.75 mm DV). Once positioned, 300 nL of BDA was infused at 0.125 μL/min. Following infusion, the needle was left for 2-5 minutes before being carefully retracted. The skin was then glued back together and the mouse gently removed from the facemask and stereotaxic frame and allowed to recover on a heating mat before being returned to the home cage.

#### BDA tissue collection and processing (*Extended Data Fig. 9I-J*)

For quantification of BDA-labeled axonal projections through the corpus callosum, animals were deeply anaesthetized (i.p. pentobarbital 100 mg/kg) and transcardially perfused with saline followed by 4% PFA 1 week after tracer injection (14-days post-stroke). Brains were dissected free, post-fixed and 40 μm thick coronal sections collected into 24-well plates containing cryoprotectant. Tissue was stored at -20 °C until required. To visualize BDA fiber staining, sections were incubated with avidin–biotin–peroxidase complex (Vectastain) followed by diaminobenzidine (DAB). Sections were mounted onto gelatin-coated glass slides, lightly airdried, passed sequentially through alcohols (50%, 70%, 95% and 100%) before being passed through xylene and then coverslipped using DPX mounting solution before being processed for densitometric analysis of axonal labels.

#### Temperature recordings in mice (*Extended Data Fig. 9A*)

Ethical permission for the following procedures were granted by the Danish Animal Experiments Inspectorate (permission 2017-15-0201-01248), and all animal procedures were performed in compliance with Directive 2010/63/EU of the European Parliament and of the Council, and with Danish Law and Order regulating animal experiments (LBK no. 253, 08/03/2013 and BEK no. 88, 30/01/2013). Mice (C57BL/6*J*Rj, Janvier, 8 weeks, *n* = 8) were habituated with i.p. injections (0.9% saline) for 4 days prior to the experiment to minimize stress on the day of the experiment. Experiments were conducted in a quiet room (22-23 °C), in which mice were left undisturbed for at least 2 h prior to the experiment. The core body temperature was measured rectally by a thermometer (Harvard Apparatus, Edenbridge, UK) via a lubricated thermistor probe (1.6 mm diameter OD probe; Harvard Apparatus) at various time points after drug or vehicle administration. Mice were held at the base of the tail and measured until a stable temperature was obtained (approx. 15 sec). Data are means ± SD, and statistical analysis used is two-way ANOVA followed by Dunnett’s post-hoc test with time and treatment as factors.

### Data availability

All raw mass spectrometry proteomics data from this study have been deposited to the ProteomeXchange Consortium via the PRIDE partner repository^58^, with the dataset identifier PXD019679. Reviewer account details: Username, Password: hgtvSBPV. The atomic coordinates have been prepared for submission to the PDB and will be released upon publication. All other data are included in the paper and Supplementary Data.

## References

1. De Koninck, P. & Schulman, H. Sensitivity of CaM kinase II to the frequency of Ca^2+^ oscillations. Science 279, 227–230, doi:10.1126/science.279.5348.227 (1998).

2. Bayer, K. U. & Schulman, H. CaM Kinase: Still Inspiring at 40. Neuron 103, 380–394, doi:10.1016/j.neuron.2019.05.033 (2019).

3. Hoelz, A., Nairn, A. C. & Kuriyan, J. Crystal structure of a tetradecameric assembly of the association domain of Ca^2+^/calmodulin-dependent kinase II. Mol Cell 11, 1241–1251, doi:doi: 10.1016/s1097-2765(03)00171-0 (2003).

4. Chao, L. H. et al. A mechanism for tunable autoinhibition in the structure of a human Ca^2+^/calmodulin-dependent kinase II holoenzyme. Cell 146, 732–745, doi:10.1016/j.cell.2011.07.038 (2011).

5. Stratton, M. et al. Activation-triggered subunit exchange between CaMKII holoenzymes facilitates the spread of kinase activity. Elife 3, e01610, doi:10.7554/eLife.01610 (2013).

6. Bhattacharyya, M. et al. Molecular mechanism of activation-triggered subunit exchange in Ca^2+^/calmodulin-dependent protein kinase II. Elife 5:e13405, doi:10.7554/eLife.13405 (2016).

7. Sloutsky, R. et al. Heterogeneity in human hippocampal CaMKII transcripts reveals allosteric hub-dependent regulation. Sci Signal 13, doi:10.1126/scisignal.aaz0240 (2020).

8. Coultrap, S. J., Vest, R. S., Ashpole, N. M., Hudmon, A. & Bayer, K. U. CaMKII in cerebral ischemia. Acta Pharmacol Sin 32, 861–872, doi:10.1038/aps.2011.68 (2011).

9. Deng, G. et al. Autonomous CaMKII activity as a drug target for histological and functional neuroprotection after resuscitation from cardiac arrest. Cell Rep 18, 1109–1117, doi:10.1016/j.celrep.2017.01.011 (2017).

10. Bernasconi, R., Mathivet, P., Bischoff, S. & Marescaux, C. Gamma-hydroxybutyric acid: an endogenous neuromodulator with abuse potential? Trends Pharmacol Sci 20, 135–141, doi:10.1016/s0165-6147(99)01341-3 (1999).

11. Rosenberg, O. S., Deindl, S., Sung, R. J., Nairn, A. C. & Kuriyan, J. Structure of the autoinhibited kinase domain of CaMKII and SAXS analysis of the holoenzyme. Cell 123, 849–860, doi:10.1016/j.cell.2005.10.029 (2005).

12. McSpadden, E. D. et al. Variation in assembly stoichiometry in non-metazoan homologs of the hub domain of Ca^2+^/calmodulin-dependent protein kinase II. Protein Sci 28, 1071–1082, doi:10.1002/pro.3614 (2019).

13. Rosenberg, O. S. et al. Oligomerization states of the association domain and the holoenyzme of Ca^2+^/CaM kinase II. FEBS J 273, 682–694, doi:10.1111/j.1742-4658.2005.05088.x (2006).

14. Torres-Ocampo, A. P. et al. Characterization of CaMKIIalpha holoenzyme stability. Protein Sci 29, 1524–1534, doi:10.1002/pro.3869 (2020).

15. Chia, P. H. et al. A homozygous loss-of-function CAMK2A mutation causes growth delay, frequent seizures and severe intellectual disability. Elife 7:e32451, doi:10.7554/eLife.32451 (2018).

16. Skelding, K. A., Spratt, N. J., Fluechter, L., Dickson, P. W. & Rostas, J. A. alphaCaMKII is differentially regulated in brain regions that exhibit differing sensitivities to ischemia and excitotoxicity. J Cereb Blood Flow Metab 32, 2181–2192, doi:10.1038/jcbfm.2012.124 (2012).

17. Ahmed, M. E. et al. Beneficial effects of a CaMKIIa inhibitor TatCN21 peptide in global cerebral ischemia. J Mol Neurosci 61, 42–51, doi:10.1007/s12031-016-0830-8 (2017).

18. Bayer, K. U., De Koninck, P., Leonard, A. S., Hell, J. W. & Schulman, H. Interaction with the NMDA receptor locks CaMKII in an active conformation. Nature 411, 801–805, doi:10.1038/35081080 (2001).

19. Vest, R. S., Davies, K. D., O’Leary, H., Port, J. D. & Bayer, K. U. Dual mechanism of a natural CaMKII inhibitor. Mol Biol Cell 18, 5024–5033, doi:10.1091/mbc.E07-02-0185 (2007).

20. Vest, R. S., O’Leary, H., Coultrap, S. J., Kindy, M. S. & Bayer, K. U. Effective post-insult neuroprotection by a novel Ca^2+^/ calmodulin-dependent protein kinase II (CaMKII) inhibitor. J Biol Chem 285, 20675–20682, doi:10.1074/jbc.M109.088617 (2010).

21. Ottani, A. et al. Effect of γ-hydroxybutyrate in two rat models of focal cerebral damage. Brain Res 986, 181–190, doi:10.1016/s0006-8993(03)03252-9 (2003).

22. Kaupmann, K. et al. Specific γ-hydroxybutyrate-binding sites but loss of pharmacological effects of γ-hydroxybutyrate in GABA_B_(1)-deficient mice. Eur J Neurosci 18, 2722–2730, doi:10.1111/j.1460-9568.2003.03013 (2003).

23. Bay, T., Eghorn, L. F., Klein, A. B. & Wellendorph, P. GHB receptor targets in the CNS: Focus on high-affinity binding sites. Biochem Pharmacol 87, 220–228, doi:10.1016/j.bcp.2013.10.028 (2014).

24. Wellendorph, P. et al. Novel cyclic γ-hydroxybutyrate (GHB) analogs with high affinity and stereoselectivity of binding to GHB sites in rat brain. J Pharmacol Exp Ther 315, 346–351, doi:10.1124/jpet.105.090472 (2005).

25. Bantscheff, M. et al. Quantitative chemical proteomics reveals mechanisms of action of clinical ABL kinase inhibitors. Nat Biotechnol 25, 1035–1044, doi:10.1038/nbt1328 (2007).

26. Baucum, A. J., 2nd, Shonesy, B. C., Rose, K. L. & Colbran, R. J. Quantitative proteomics analysis of CaMKII phosphorylation and the CaMKII interactome in the mouse forebrain. ACS Chem Neurosci 6, 615–631, doi:10.1021/cn500337u (2015).

27. Vogensen, S. B. et al. New synthesis and tritium labeling of a selective ligand for studying high-affinity γ-hydroxybutyrate (GHB) binding sites. J Med Chem 56, 8201–8205, doi:10.1021/jm4011719 (2013).

28. Ashpole, N. M. & Hudmon, A. Excitotoxic neuroprotection and vulnerability with CaMKII inhibition. Mol Cell Neurosci 46, 720–730, doi:10.1016/j.mcn.2011.02.003 (2011).

29. Buonarati, O. R. et al. CaMKII versus DAPK1 binding to GluN2B in ischemic neuronal cell death after resuscitation from cardiac arrest. Cell Rep 30, 1–8 e4, doi:10.1016/j.celrep.2019.11.076 (2020).

30. Clarkson, A. N., Huang, B. S., Macisaac, S. E., Mody, I. & Carmichael, S. T. Reducing excessive GABA-mediated tonic inhibition promotes functional recovery after stroke. Nature 468, 305–309, doi:10.1038/nature09511 (2010).

31. Singer, W. Development and plasticity of cortical processing architectures. Science 270, 758–764, doi:10.1126/science.270.5237.758 (1995).

32. Gao, B., Kilic, E., Baumann, C. R., Hermann, D. M. & Bassetti, C. L. Gammahydroxybutyrate accelerates functional recovery after focal cerebral ischemia. Cerebrovasc Dis 26, 413–419, doi:10.1159/000151683 (2008).

33. Busardo, F. P., Kyriakou, C., Napoletano, S., Marinelli, E. & Zaami, S. Clinical applications of sodium oxybate (GHB): from narcolepsy to alcohol withdrawal syndrome. Eur Rev Med Pharmacol Sci 19, 4654–4663 (2015).

34. Feigin, V. L., Norrving, B. & Mensah, G. A. Global burden of stroke. Circ Res 120, 439–448, doi:10.1161/CIRCRESAHA.116.308413 (2017).

## Additional references

35. Vogensen, S. B. et al. New synthesis and tritium labeling of a selective ligand for studying high-affinity γ-hydroxybutyrate (GHB) binding sites. J Med Chem 56, 8201–8205, doi:10.1021/jm4011719 (2013).

36. Bhattacharyya, M. et al. Molecular mechanism of activation-triggered subunit exchange in Ca^2+^/calmodulin-dependent protein kinase II. Elife 5, e13405, doi:10.7554/eLife.13405 (2016).

37. Proietti Onori, M. et al. The intellectual disability-associated CAMK2G p.Arg292Pro mutation acts as a pathogenic gain-of-function. Hum Mutat 39, 2008–2024, doi:10.1002/humu.23647 (2018).

38. McSpadden, E. D. et al. Variation in assembly stoichiometry in non-metazoan homologs of the hub domain of Ca^2+^/calmodulin-dependent protein kinase II. Protein Sci 28, 1071–1082, doi:10.1002/pro.3614 (2019).

39. Kool, M. J., van de Bree, J. E., Bodde, H. E., Elgersma, Y. & van Woerden, G. M. The molecular, temporal and region-specific requirements of the beta isoform of Calcium/Calmodulin-dependent protein kinase type 2 (CAMK2B) in mouse locomotion. Sci Rep 6, 26989, doi:10.1038/srep26989 (2016).

40. Sabbatini, P. et al. Design, synthesis and in vitro pharmacology of new radiolabelled GHB analogues including photolabile analogues with irreversible binding to the high-affinity GHB binding sites. J Med Chem 53, 6506–6510, doi: 10.1021/jm1006325 (2010).

41. Wellendorph, P. et al. Novel cyclic γ-hydroxybutyrate (GHB) analogs with high affinity and stereoselectivity of binding to GHB sites in rat brain. J Pharmacol Exp Ther 315, 346–351, doi:10.1124/jpet.105.090472 (2005).

42. Cox, J. et al. Accurate proteome-wide label-free quantification by delayed normalization and maximal peptide ratio extraction, termed MaxLFQ. Mol Cell Proteomics 13, 2513–2526, doi:10.1074/mcp.M113.031591 (2014).

43. Hoelz, A., Nairn, A. C. & Kuriyan, J. Crystal structure of a tetradecameric assembly of the association domain of Ca^2+^/calmodulin-dependent kinase II. Mol Cell 11, 1241–1251, doi:doi: 10.1016/s1097-2765(03)00171-0 (2003).

44. Kabsch, W. Xds. Acta Crystallogr D Biol Crystallogr 66, 125–132, doi:10.1107/S0907444909047337 (2010).

45. Evans, P. R. & Murshudov, G. N. How good are my data and what is the resolution? Acta Crystallogr D Biol Crystallogr 69, 1204–1214, doi:10.1107/S0907444913000061 (2013).

46. Winn, M. D. et al. Overview of the CCP4 suite and current developments. Acta Crystallogr D Biol Crystallogr 67, 235–242, doi:10.1107/S0907444910045749 (2011).

47. Adams, P. D. et al. PHENIX: a comprehensive Python-based system for macromolecular structure solution. Acta Crystallogr D Biol Crystallogr 66, 213–221, doi:10.1107/S0907444909052925 (2010).

48. Emsley, P., Lohkamp, B., Scott, W. G. & Cowtan, K. Features and development of Coot. Acta Crystallogr D Biol Crystallogr 66, 486–501, doi:10.1107/S0907444910007493 (2010).

49. Moriarty, N. W., Grosse-Kunstleve, R. W. & Adams, P. D. electronic Ligand Builder and Optimization Workbench (eLBOW): a tool for ligand coordinate and restraint generation. Acta Crystallogr D Biol Crystallogr 65, 1074–1080, doi:10.1107/S0907444909029436 (2009).

50. Morin, A. et al. Collaboration gets the most out of software. Elife 2, e01456, doi:10.7554/eLife.01456 (2013).

51. Sastry, G. M., Adzhigirey, M., Day, T., Annabhimoju, R. & Sherman, W. Protein and ligand preparation: parameters, protocols, and influence on virtual screening enrichments. J Comput Aided Mol Des 27, 221–234, doi:10.1007/s10822-013-9644-8 (2013).

52. Shelley, J. C. et al. Epik: a software program for pK(a) prediction and protonation state generation for drug-like molecules. J Comput Aided Mol Des 21, 681–691, doi:10.1007/s10822-007-9133-z (2007).

53. Banker, G. & Goslin, K. Culturing Nerve Cells. (MIT Press, 1991).

54. Vest, R. S., O’Leary, H., Coultrap, S. J., Kindy, M. S. & Bayer, K. U. Effective post-insult neuroprotection by a novel Ca^2+^/ calmodulin-dependent protein kinase II (CaMKII) inhibitor. J Biol Chem 285, 20675–20682, doi:10.1074/jbc.M109.088617 (2010).

55. Overman, J. J. et al. A role for ephrin-A5 in axonal sprouting, recovery, and activity-dependent plasticity after stroke. Proc Natl Acad Sci USA 109, E2230–2239, doi:10.1073/pnas.1204386109 (2012).

56. Clarkson, A. N., Huang, B. S., Macisaac, S. E., Mody, I. & Carmichael, S. T. Reducing excessive GABA-mediated tonic inhibition promotes functional recovery after stroke. Nature 468, 305–309, doi:10.1038/nature09511 (2010).

57. Crawford, D. K., Mangiardi, M., Xia, X., Lopez-Valdes, H. E. & Tiwari-Woodruff, S. K. Functional recovery of callosal axons following demyelination: a critical window. Neuroscience 164, 1407–1421, doi:10.1016/j.neuroscience.2009.09.069 (2009).

58. Vizcaino, J. A. et al. ProteomeXchange provides globally coordinated proteomics data submission and dissemination. Nat Biotechnol 32, 223–226, doi:10.1038/nbt.2839 (2014).

