## Supplementary figures for "GHB confers neuroprotection by stabilizing the CaMKIIα hub domain"

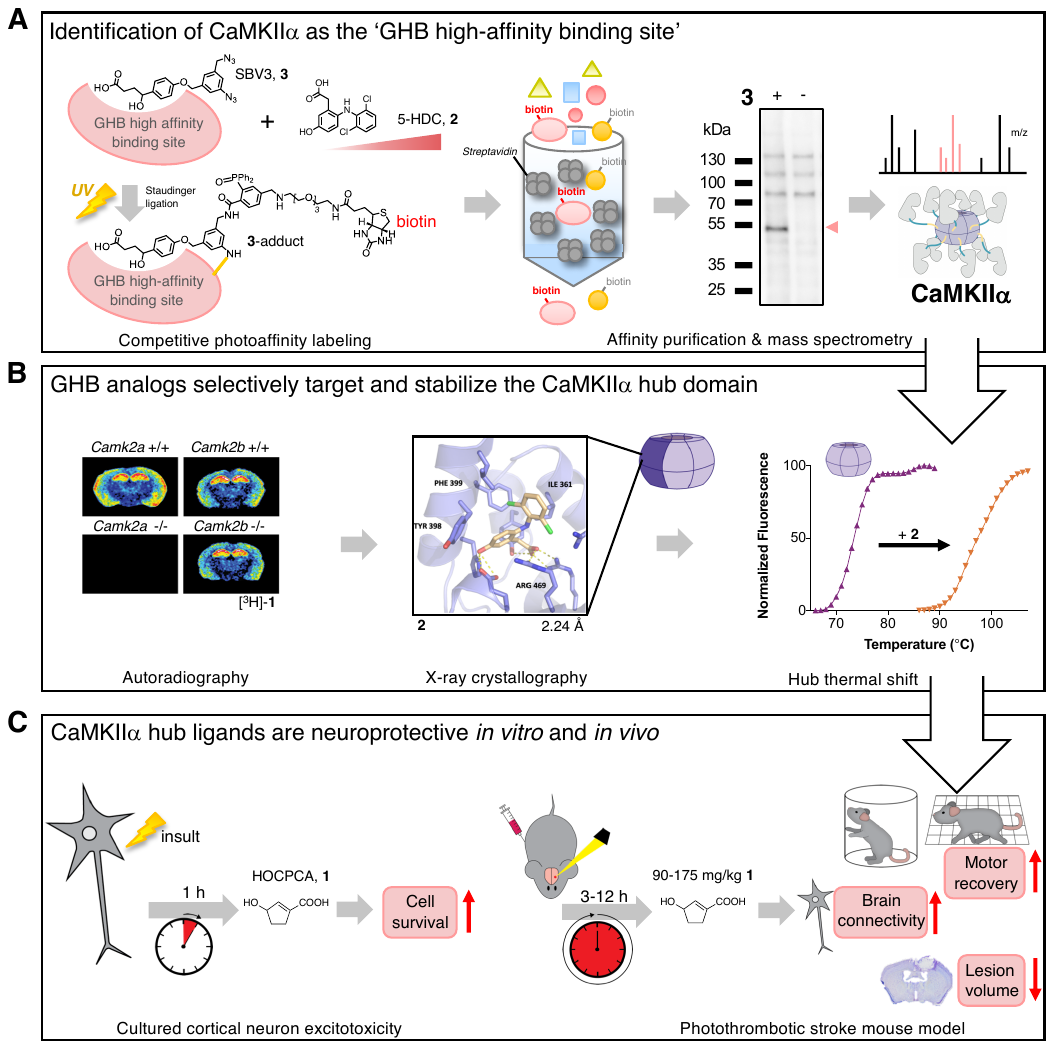

**Extended Data Fig. 1. Schematic summary of key results.** (A) CaMKIIα was identified as the ‘GHB high-affinity binding site’ with a workflow consisting of competitive photoaffinity labeling (PAL), affinity purification (AP) and quantitative chemical proteomics. Rat hippocampal homogenate was labeled with the photoligand **3**, containing a GHB-binding moiety, an aryl azide group for photolabeling, and an alkyl azide group for subsequent ligation of a phosphine-PEG_3_-biotin linker. PAL was competed with free **2**. After streptavidin AP of biotinylated proteins, the enriched binding protein was subjected to SDS-PAGE, in-gel digestion and LC-MS/MS analysis. (B) The clear absence of [^3^H]-**1** binding to *Camk2a* knockout (-/-) mouse brain tissue validates CaMKIIα as the GHB binding protein. A deep binding cavity in the hub domain was identified as the binding site for the ligand **2** by co-crystallization. Compound **2** was shown to dramatically stabilize the hub in a thermal shift assay. (C) **1** shows neuroprotective effects both *in vitro* in cultured cortical neurons after Glu exposure and *in vivo* in mice when administered up to 12 h after a photothrombotic stroke and assessed 1 week later.

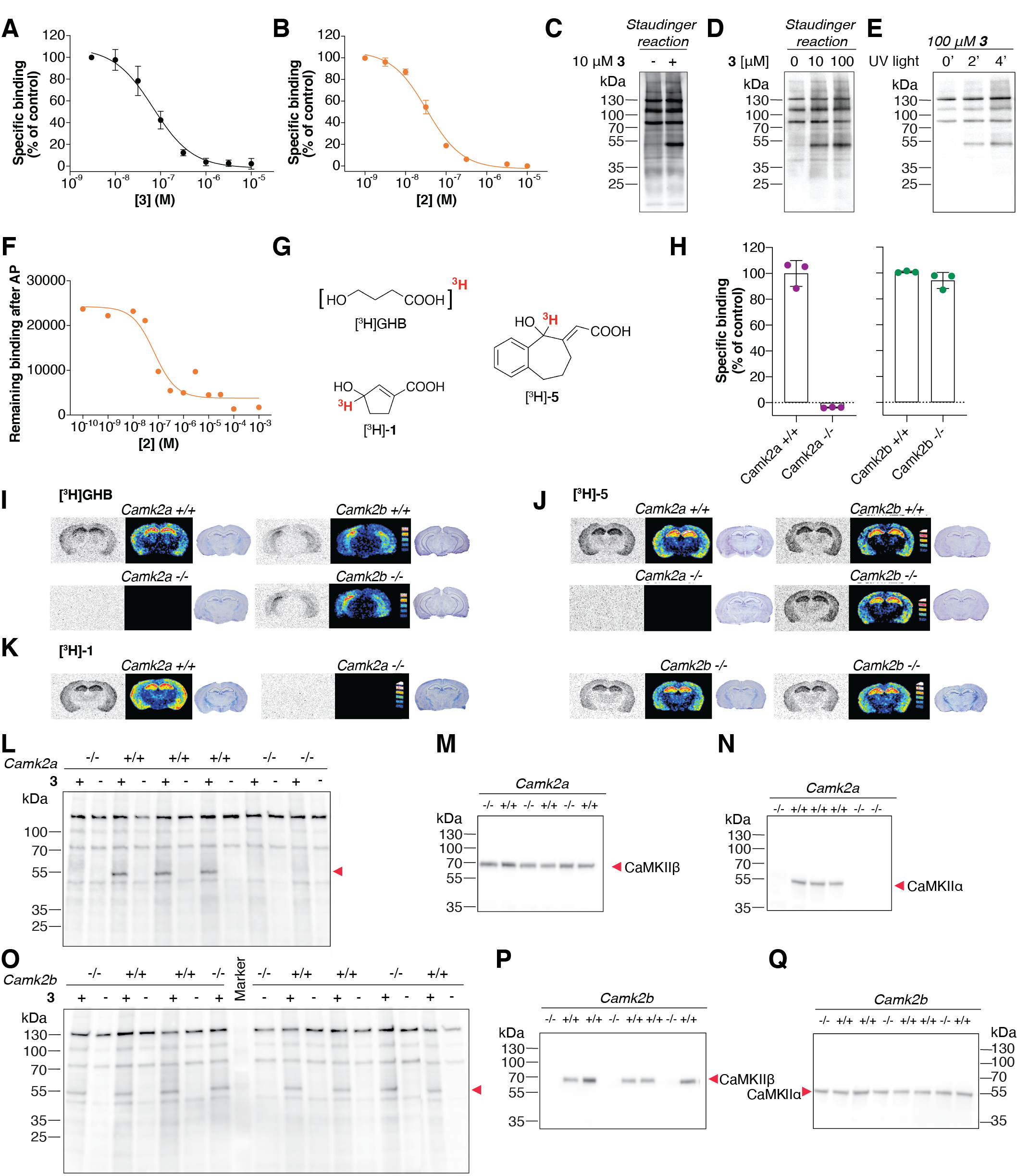

Extended Data Fig. 2. Validation of the CaMKIIα binding site. (A-F) Photolabeling with 3 to rat cortical homogenate. (A, B) Concentration-dependent inhibition of [^3^H]NCS-382 binding by 3, and 2. Results are given as means ± SEM of (*n* = 3-5). Average *K*_I_ (p*K*_I_ ± SEM) values were 66 nM (7.2 ± 0.104) and 35 nM (7.5 ± 0.05), respectively. (C) Optimization of photolabeling (shown with biotinylation) achieved by Staudinger-Bertozzi ligation followed by anti-biotin Western blotting. The signature band at ~55 kDa is detectable only after photolinking with SBV3 (10 μM). (D) Optimization of SBV3 concentration prior to photolabeling. (E) Optimization of UV irradiation exposure time (in min) for photolabeling with SBV3. (F) Concentration-dependent inhibition of the SBV3-induced PAL-AP reaction by 2. IC_50_ value obtained 69 nM (single experiment). (G) Chemical structures of radioligands. (H) No [^3^H]-1(5 nM) binding to cortical homogenate from *Camk2a* -/- cf. *Camk2a* +/+, *Camk2b* -/- and *Camk2a* +/+ samples (unpaired t-test with Welch’s correction (*n* = 3). (I-K) *In vitro* autoradiography with [^3^H]GHB (30 nM), [^3^H]NCS-382 (7 nM), and [^3^H]-1 (1 nM) confirms absence of binding only in *Camk2a* -/- tissue. Representative autoradiograms are supported by pseudo color images as well as cresyl violet staining of coronal section. (*n* = 3-4, 4 sections per animal). (L) Anti-biotin Western blots showing the complete absence of the 55 kDa-photoaffinity-labeled band in *Camk2a* -/- cf. +/+ samples. (M) and (N) Anti-CaMKIIα and CaMKIIβ Western blots of *Camk2a* -/- cf. +/+ brain samples, confirming the deletion of CaMKIIα only. (O) Anti-biotin Western blot showing an intact 55 kDa band in both *Camk2b* +/+ and -/- samples. (P, Q) Anti-CaMKIIα and CaMKIIβ Western blots of *Camk2b* -/- cf. +/+ brain samples, confirming the deletion of CaMKIIβ only. (L-Q) Red arrows indicate relevant bands.

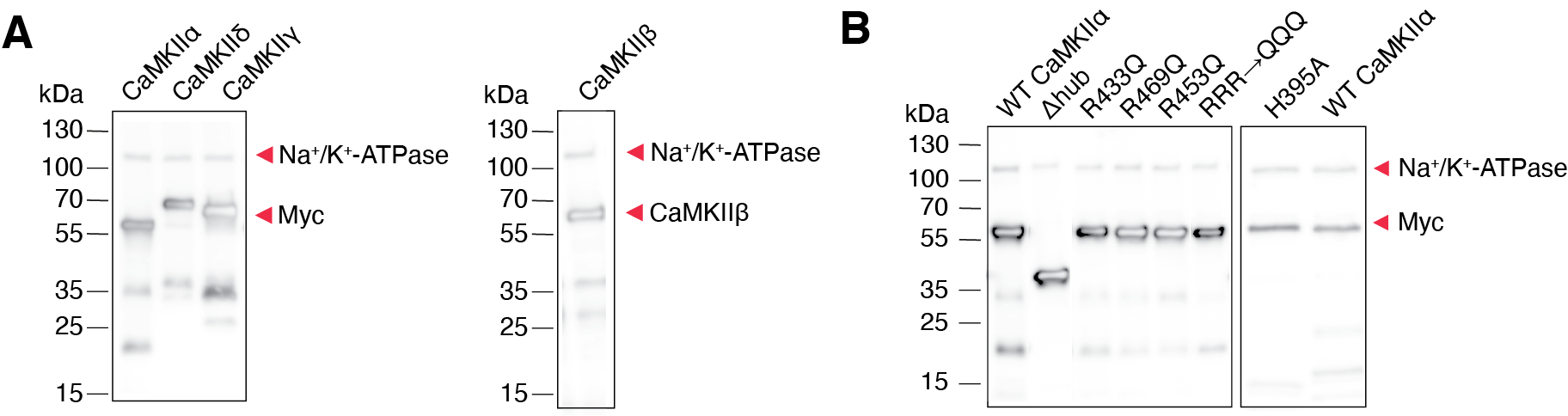

Extended Data Fig. 3. Western blot validation of myc-tagged CaMKIIα/γ/δ, CaMKIIβ expression or CaMKIIα mutants in cell homogenate from transiently expressing HEK293T cells. (A) Red arrows indicate the relevant bands at 55 kDa for CaMKIIα, 70 kDa for CaMKIIδ, 65 kDa for CaMKIIγ and 60 kDa for CaMKIIβ corresponding to the expected sizes of the subtypes plus the c-terminal myc tag (not present in CaMKIIβ). Bands for Na^+^/K^+^-ATPase at 110 kDa confirmed equal loading of samples. (B) Red arrows indicate the relevant bands at around 55 kDa corresponding to the expected size of CaMKIIα-myc. The mutant lacking the hub domain (Δhub) shows a band located at 35 kDa corresponding to a smaller protein. Bands for Na^+^/K^+^-ATPase at 110 kDa confirmed equal loading of samples.

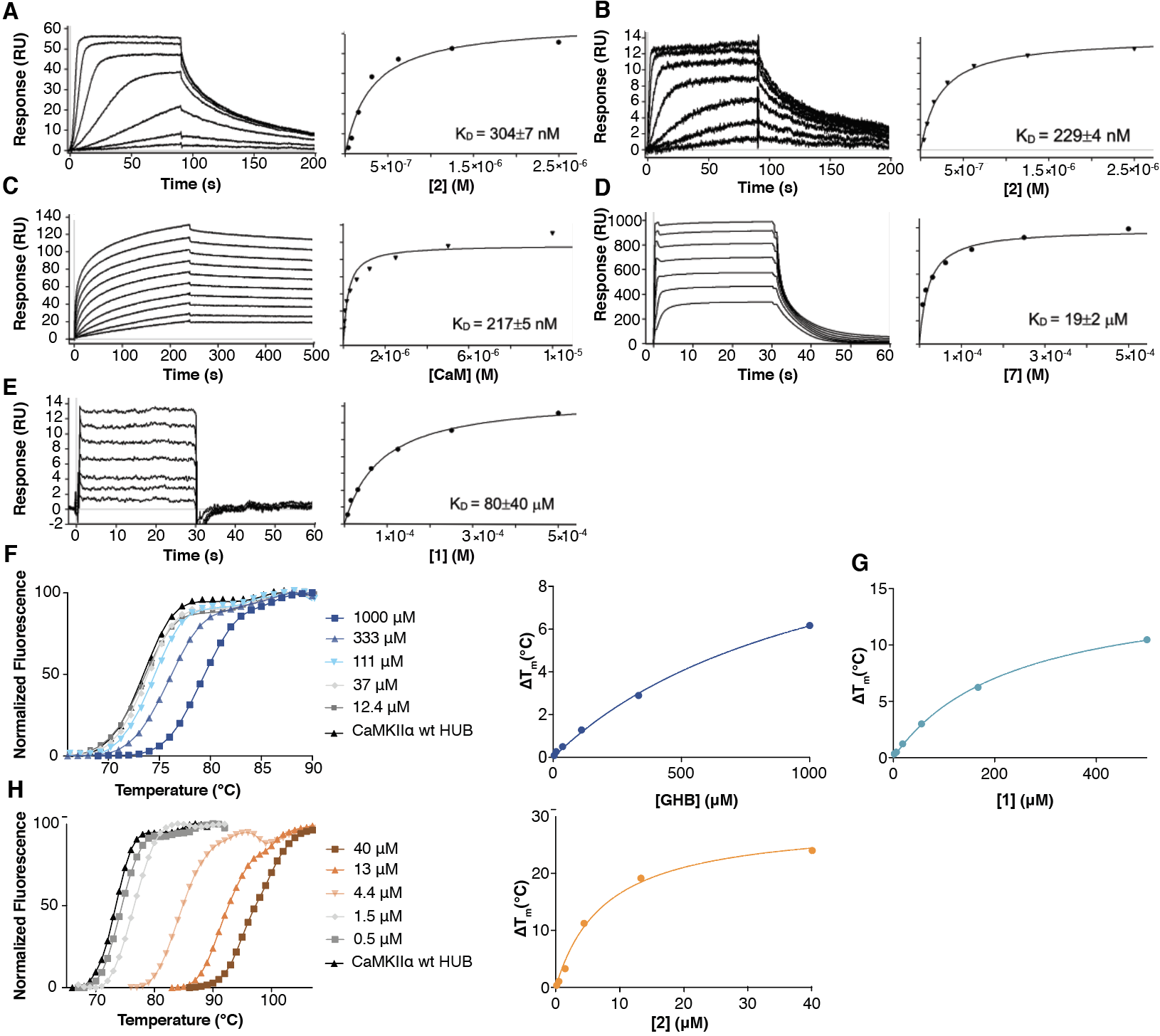

**Extented Data Fig. 4. Biophysical experiments with CaMKIIα hub ligands**. (A-E) SPR sensorgrams of ligands interacting with immobilized CaMKIIα. Compounds were injected in 2-fold serial dilutions over immobilized CaMKIIα (WT hub, 6X Hub or full-length, as specified). (A) Binding of **2** is unchanged to CaMKIIα WT hub (cf. 6x Hub in Fig. 2B), and (B) full-length CaMKIIα. (C) CaM control binding to immobilized full-length CaMKIIα (pH 7.4). The binding of CaM to CaMKIIα full-length was fully regenerated between concentrations by injection of 1 mM EDTA. (D) Control peptide **7** and (E) binding of **1** to the 6x Hub (pH 6 as for **2** in Fig. 2B). SPR sensorgrams are blank injection and reference surface subtracted (*left* panels). Plots of equilibrium binding responses at the end of the analyte injections against analyte concentration (*right* panels). Steady state *K*_D_ ± SEM were derived from curves fitted to a 1:1 model based on at least seven concentration-response measurements (Extended Table 1c). (F-H) Melting curves from differential scanning fluorimetry of the CaMKIIα WT hub upon GHB, **1** or **2** binding. TSA melting curves (*left* panels) and ∆T_m_ of CaMKIIα hub by compounds, derived from plots of T_m_ against analyte concentration fitted to a 1:1 model (*right* panels) of (F) GHB (12-1000 μM), (G) **1** (6-500 μM); melting curve is given in main text, and (H) **2** (0.5-40 μM). The ∆T_m_ of CaMKIIα by GHB, **1** and **2** were estimated to 13.0 °C, 15.3 °C and 29.2 °C, respectively.

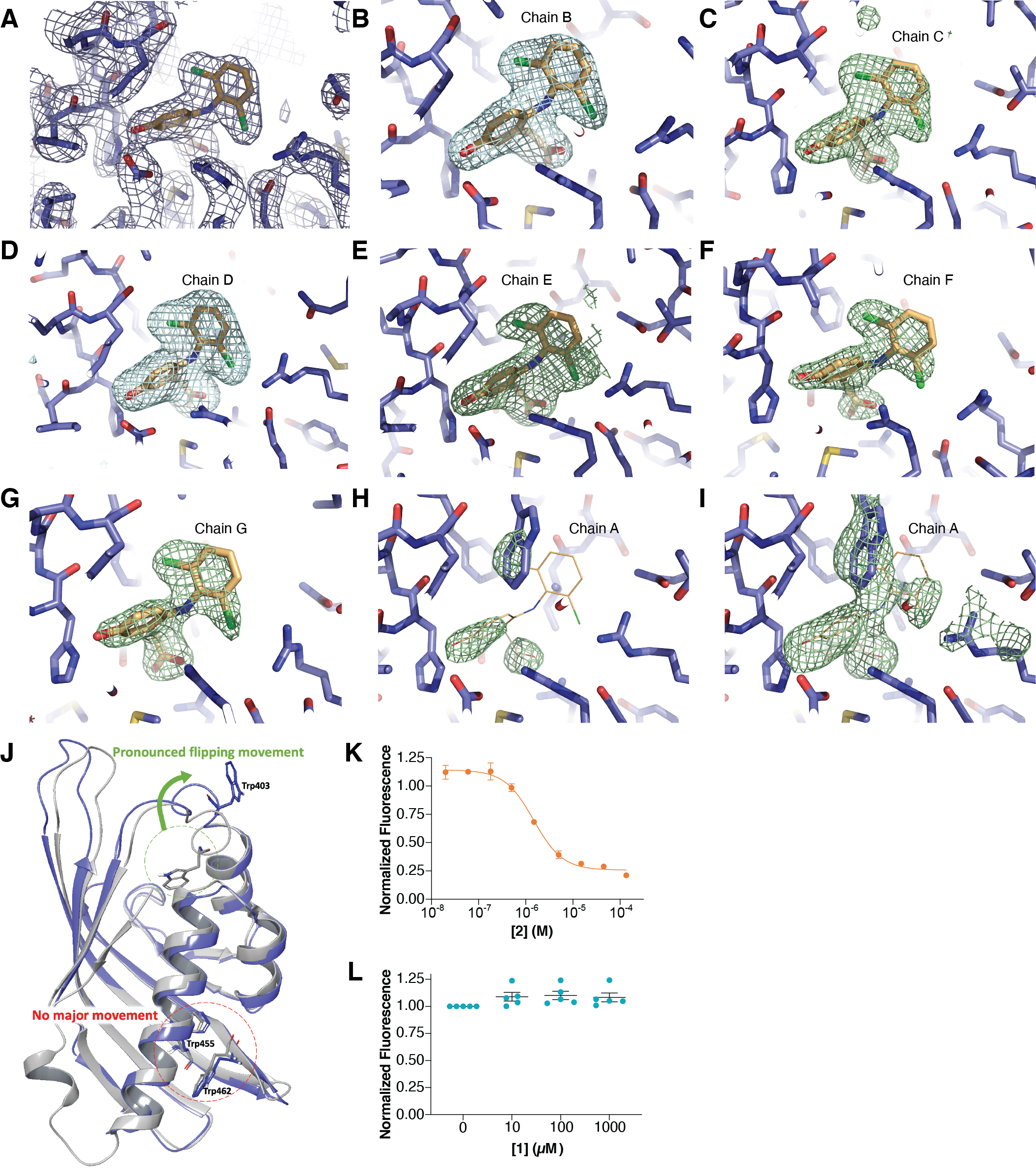

**Extended Data Fig. 5. Data to support structural findings.** (A-I) Electron density map of **2** bound to CaMKIIα 6x Hub. (A) 2Fo-Fc density showing **2** bound to a pocket of the hub monomer. (B-G) Polder omit maps (Ref 10) where the ligand in the chain indicated is omitted in the map calculation, along with the surrounding solvent. Maps are contoured at 4.5 sigma. (H) Polder omit map for the pocket with the tryptophan (Trp) modeled inwards. Trp403 was omitted along with the nearby ordered waters and the surrounding solvent. The map was contoured at 4.5 sigma and where the ligand would be expected to bind it is shown with a line model (this ligand is not modeled in the structure). (I) Same as in (H) but the map is contoured at 3 sigma revealing weak density for the ligand as well as Trp403 and therefore a mixture of the two states in the crystal. (J) Protein superimposition comparing the relative positioning of the three hub domain Trp residues without (grey, PDB entry 5IG3) and with **2** bound (purple). Upon ligand binding there is a pronounced flip of Trp403 but no movement of Trp455 and Trp462. (Concentration-dependent quenching of Trp403 fluorescence with (K) Compound **2** (IC_50_ value of 1.47 μM) and (L) **1** (no change). For **2**, the graph is based on pooled data (normalized means ± SEM) of two technical replicates. For **1**, each data point represents independent data as means of five technical replicates.

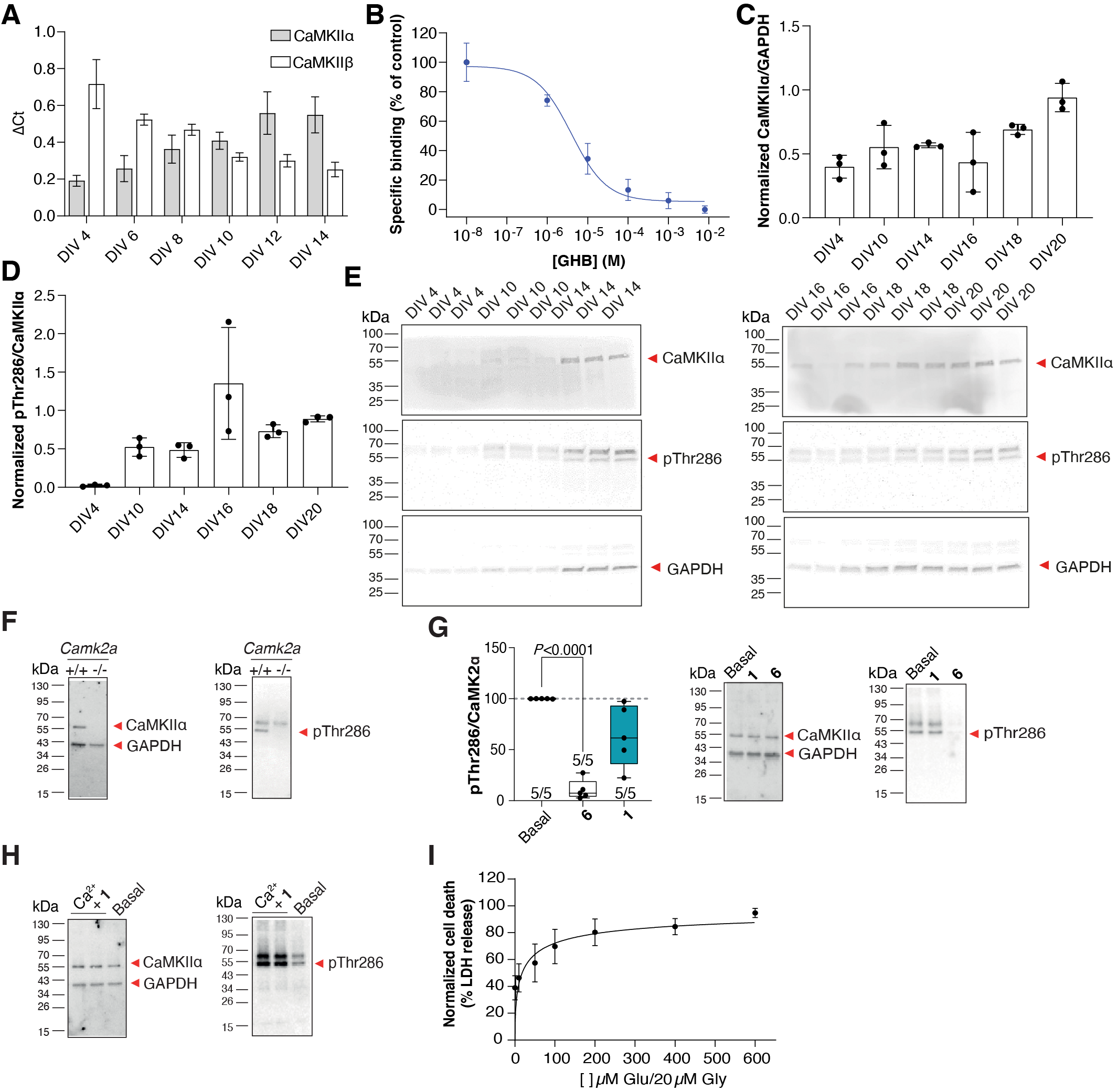

**Extended Data Fig. 6. Supporting data for pharmacological data on cortical neuronal cultures.** (A-E) Optimization of neuronal culturing. (A) mRNA levels of *Camk2a* increase during culturing time (grey bars) whereas *Camk2b* decreases (white bars). Data is presented as means ± SEM of three individual cultures in technical triplicates. (B) Concentration-dependent inhibition of [^3^H]-**1** binding to neuronal culture cell homogenate by GHB. Shown is representative data ± SD of technical triplicates. The average IC_50_ value (pIC_50_ ± SEM) obtained was 2.04 μM (5.7 ± 0.07) (three different cultures). Western blots show a temporal increase in protein levels of CaMKIIα (C) and pThr286 (D). Data is represented as means ± SD from three different wells from the same culture. (E) Corresponding representative western blots. (F) Representative Western blots showing the expected absence of CaMKIIα (*left*) and pThr286 bands (*right*) in cortical neuronal cultures from *Camk2a* -/- cf. +/+ cultures. Data is represented as means ± SD from three different wells from the same culture. (G) Effects of **1** (3 mM) and KN93 (20 μM) on basal pThr286 levels in cortical neuronal cultures and corresponding representative Western blots (Brown-Forsythe and Welch ANOVA, post-hoc Dunnett’s T3 test). Number in bar diagrams indicates number of experiments/individual cultures. Box plots (boxes, 25–75%; whiskers, minimum and maximum; lines, median). (H) Representative Western blots of effect of **1** (3 mM) on Ca^2+^-stimulated (100 μM) pThr286 levels. *For G:* (I) Neuronal cell death determined with LDH release after an excitotoxic insult. Maximum cell death was obtained with 100 μM Glu/20 μM glycine and above. Results are normalized to maximum cell death corresponding to the highest Glu concentration. Pooled data (mean ± SEM) (five different cultures, technical triplicates).

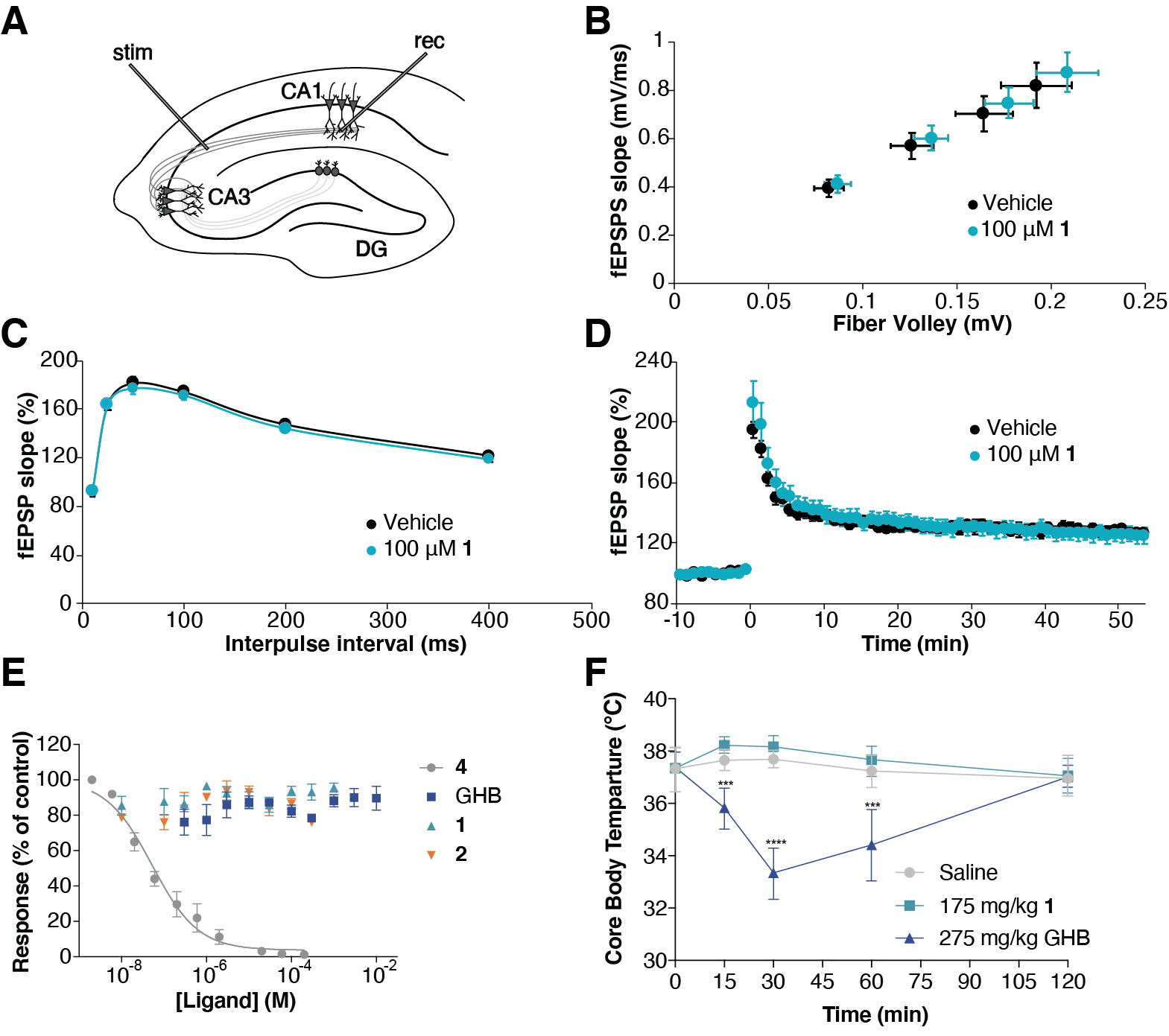

**Extended Data Fig. 7. Supporting data showing lack of effects of compound 1.** (A-D) Lack of effect in long-term potentiation (LTP). (A) Schematic overview of LTP induction in the CA3–CA1 pathway. stim, stimulating electrode; rec, recording electrode; DG, dentate gyrus. (B) Basal synaptic transmission is unaffected upon influx of **1** (100 μM) [vehicle (*n* = 32 from 6 mice), **1** (*n* = 24 from five mice)]. (C) Paired-pulse facilitation (PPF) is unaffected upon influx of **1** (100 μM) [vehicle (*n* = 28 from 6 mice), **1** (*n* = 20 from 4 mice)]. (D) LTP is unaffected upon influx of **1** (100 μM) [vehicle (*n* = 23 from 6 mice), **1** (*n* = 22 from five mice)]. (Mean ± SEM, Repeated Measures ANOVA). (E) Effects of GHB and analogs on CaMKIIα substrate phosphorylation using syntide-II (ADP-Glo). CaMKIIα full-length protein was used for the experiments. Compound **4** robustly inhibits syntide-II phosphorylation. GHB, **1** and **2** are inactive. Reactions were incubated for 60 min at 37 °C. Kinase reactions without enzyme or substrate were used as controls. Shown are pooled data (*n* = 3, means ± SEM) each performed in technical triplicates. (F) Compound **1** does not evoke hypothermia underlining a lack of GABA_B_ receptor activity. 275 mg/kg GHB induces a reduction in core body temperature, whereas 175 mg/kg **1** had no effect compared to saline-treated control group (means ± SD, *n* = 8, two-way ANOVA with time and treatment as factors followed by Dunnett’s test).

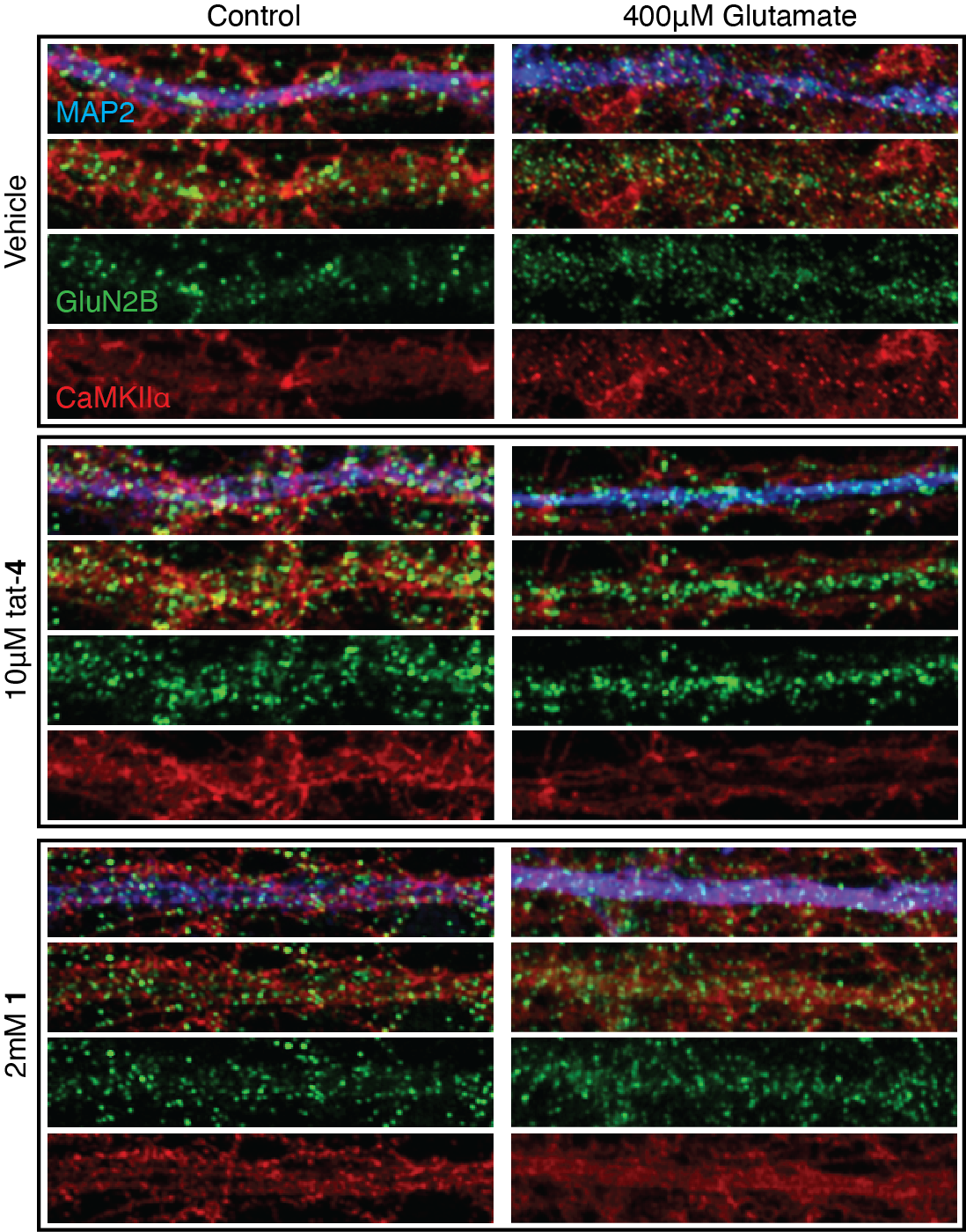

**Extended Data Fig. 8. Effect of 1 on GluN2B-CaMKIIα co-localization*.*** Representative immunostained images showing GluN2B-CaMKIIα co-localization in dendrites of hippocampal neurons (DIV 14-19) exposed to Glu 400 μM for 2 min and immediately fixed (GluN2B in green, CaMKIIα in red and MAP2 in blue). GluN2B-CaMKIIα co-localization upon stimulation was prevented by **1**, similar to tat-**4**. NB, note the dotted CaMKIIα pattern in the dendrite upon stimulation in the vehicle condition, which is absent in the **1** and tat-**4** condition.

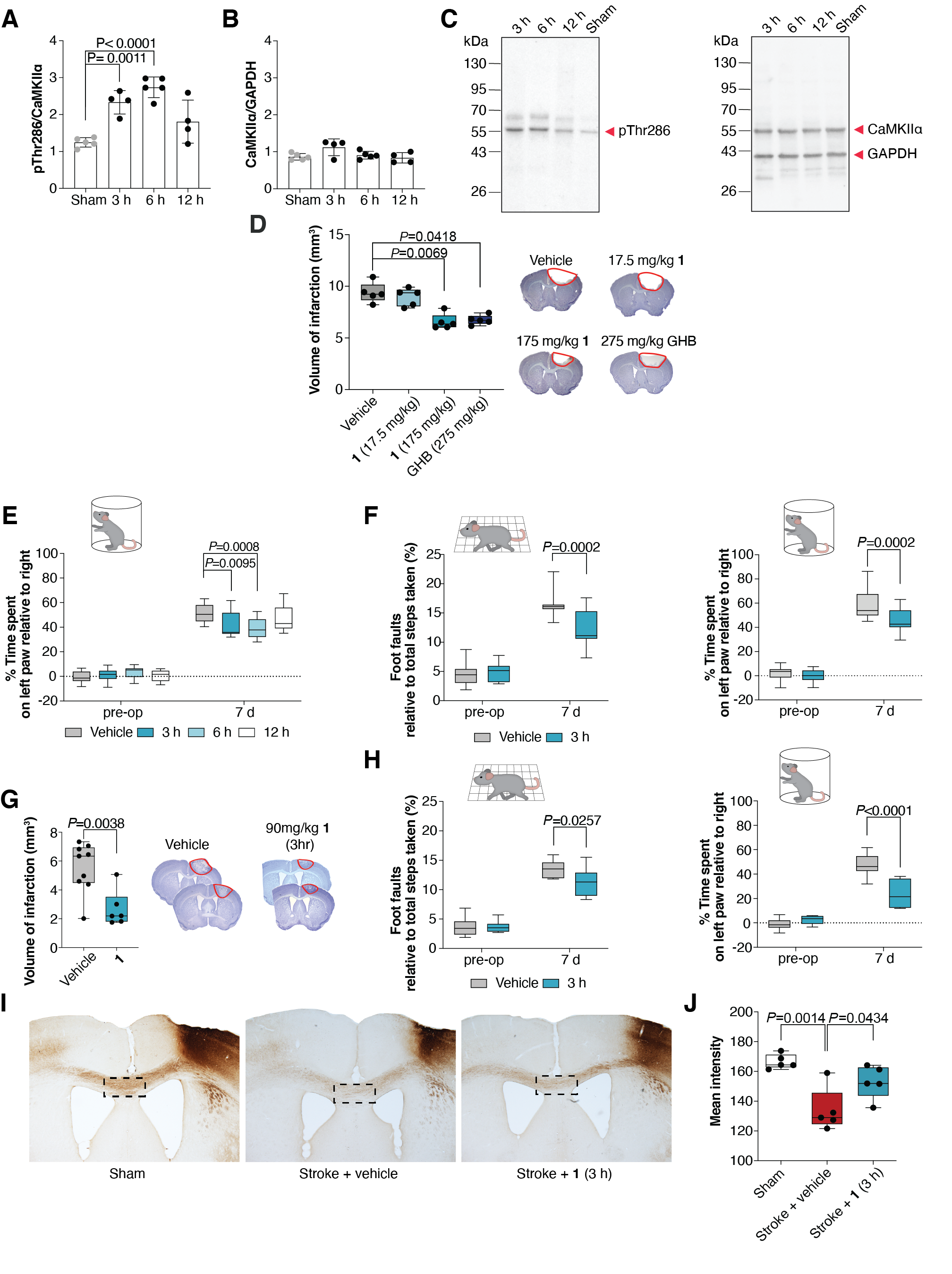

**Extended Data Fig. 9. Supporting data for *in vivo* data.** (A-C) CaMKIIα autophosphorylation and total CaMKIIα expression at different time points after photothrombosis compared to sham-operated animals. Quantification of immunoblots of (A) pThr286 levels normalized to total CaMKIIα expression and (B) total CaMKIIα expression normalized to GAPDH. (C) Representative Western blots. (*n* = 4-5, three technical repetitions, mean ± SD, One-way ANOVA, post-hoc Dunnett’s test). (D) Neuroprotective effects of a single i.p. dose of 17.5 mg/kg **1** and 275 kg/kg GHB administered 30 min post-photothrombotic stroke cf. vehicle (*n* = 5/group). Quantification of the stroke volume (*left*) and representative cresyl violet stainings of brain slices (*right*) measured 3 days post-stroke. (One-way ANOVA, post-hoc Kruskal-Wallis test; boxes, 25–75%; whiskers, minimum and maximum; lines, median). (E-F) Motor function improvement after GHB analog treatment in stroked mice. Recovery of forelimb motor function was assessed by analysis of foot faults in the grid-walking task as well as cylinder task for forelimb asymmetry pre-op (7 days before) and 7 days after photothrombotic stroke. (E) Cylinder task for male mice treated with a single i.p. dose of **1** (175 mg/kg) at 3 h (*n* = 13), 6 h (*n* = 10) or 12 h (*n* = 12) post-photothrombotic stroke cf. vehicle (*n* = 10). (F) Grid-walking (*left*) and cylinder task (*right*) for aged female mice (20-24 months) treated with single i.p. dose of **1** (175 mg/kg) at 3 h post-photothrombotic stroke (*n* = 13) cf. vehicle (*n* = 14). (Two-way ANOVA, post-hoc Dunnett’s test with time and treatment as independent factors and time as repeated measures; boxes, 25–75%; whiskers, minimum and maximum; lines, median). (G-H) Neuroprotective effects are also seen with **1** (90 mg/kg). Neuroprotective effect of a single i.p. dose of **1** (90 mg/kg) (*n* = 9) at 3 h post-photothrombotic stroke cf. vehicle (*n* = 6). (G) Quantification of the stroke volume (*left*) and representative cresyl violet stainings of brain slices (*right*) measured 7 days post-stroke. (Two-tailed Student’s *t*-test). (H) Recovery of motor function was assessed by analysis of foot faults in the grid-walking task (*left*) and cylinder task for forelimb asymmetry (*right*) pre-op (7 days before) and 7 days after stroke. (Two-way ANOVA, post-hoc Dunnett’s test with time and treatment as independent factors and time as repeated measures; boxes, 25–75%; whiskers, minimum and maximum; lines, median). (I-J) Compound **1** treatments partially prevents the loss of axonal connections through the corpus callosum. (I) Images of biotinylated dextran amine (BDA)-labeled connections to the premotor cortex for sham (*left*), Stroke + vehicle (mid) and Stroke + **1** (175 mg/kg i.p.) (*right*) treatments. (J) Quantification of axonal projections through the corpus callosum. Dotted rectangle represents region of analysis. One-way ANOVA, ** = *P*<0.01, for sham compared to stroke + vehicle; + = *P*<0.05, for stroke + **1** compared to stroke + vehicle. Box plots for n=5 per treatment group (boxes, 25–75%; whiskers, minimum and maximum; lines, median). (One-way ANOVA, post hoc Tukey’s test).

Extended Data Table 1a.

**List of best hit proteins from non-linear regression analysis (R^2^ > 0.6) identified from photoaffinity-labeling of ‘GHB high-affinity binding sites’ in rat hippocampal homogenate (*related to Fig. 1 and further described in Supplementary Dataset 1*).** Note that the signature photolabeled, biotin-ligated band fits only convincingly with CaMKIIα at ~55 kDa.

| Protein Name | Gene Name | UniProt ID | Molecular weight (kDa) | MS/MS counts | Non-linear regression analysis | |
| --- | --- | --- | --- | --- | --- | --- |
|  |  |  |  |  | **Top plateau** | **R^2^ value** |
| Calcium/calmodulin-dependent protein kinase II alpha | *Camk2a* | P11275 | 54.115 | 800 | 7.97E+09 | 0.81 |
| Tubulin alpha 1B | *Tuba1b* | Q6P9V9 | 50.152 | 680 | 2.14E+09 | 0.72 |
| Sodium/potassium-transporting ATPase beta-1 | *Atp1b1* | P07340 | 35.202 | 446 | 1.69E+09 | 0.68 |
| Tubulin beta 4B | *Tubb4b* | Q6P9T8 | 49.801 | 1066 | 1.50E+09 | 0.76 |
| Calcium/calmodulin-dependent protein kinase II beta | *Camk2b* | P08413 | 60.402 | 474 | 9.76E+08 | 0.65 |
| Calcium/calmodulin-dependent protein kinase type II subunit delta | *CaMKIId* | P15791 | 60.081 | 83 | 1.08E+08 | 0.82 |
| Calcium/calmodulin-dependent protein kinase II gamma | *CaMKIIg* | P11730 | 59.038 | 86 | 9.09E+07 | 0.63 |

**Extended Data Table 1b.**

**Saturation data of [^3^H]-1 binding to CaMKIIα**

|  | Rat cortical homogenate | rCaMKIIα-HEK |
| --- | --- | --- |
| *K*_D_ (μM) (p*K*_D_ ± SEM) | 0.26 (6.6 ± 0.06) | 1.8 (5.8 ± 0.10) |
| B_max_  (pmol/mg protein) | 43 | 64 |

Data are based on a number of independent experiments each performed in technical triplicates

(*n* = 3 for cortical homogenate, *n* = 5 for HEK293T cells). r, rat.

Assays were performed at pH 6 (*further described in methods section*).

**Extended Data Table 1c.**

**Collected inhibitory affinity constants from native and recombinant CaMKIIα binding assays**

|  | [^3^H]-1 binding | | SPR |
| --- | --- | --- | --- |
|  | **Rat cortical homogenate** | **CaMKIIα-HEK** | **6x Hub** |
|  | *K*_I_ (μM)  (pK_I_ ± SEM) (*n*) | | *K*_D_ (μM)  (p*K*_D_ ± SEM) (*n*) |
| GHB | 3.0  (5.5 ± 0.10) (4) | 51  (4.3 ± 0.05) (3) | n.d. |
| 1 | 0.13  (6.9 ± 0.07) (4) | 1.8  (5.8 ± 0.07) (7) | 58  (4.35 ± 0.25) (3) |
| 2 | 0.022  (7.7 ± 0.05) (3) | 0.80  (6.1 ± 0.05) (3) | 0.30  (6.53 ± 0.07) (3) |

*K*_I_ values were determined by means of the Cheng-Prusoff equation. Numbers in parentheses refer to the number of independent experiments each carried out in technical triplicates. n.d. *not determined*.

All assays were performed at pH 6 (*further described in methods section*).
